# Predicting correct serotypes using machine-learning models based on codon usage patterns of influenza A viruses

**DOI:** 10.1101/528083

## Abstract

Consistent codon usage patterns across species was supposed to be observed owing to the degeneracy of genetic code and the conservation of the translation machinery. In fact, however, codon usage vary dramatically among organisms, and the choice difference might also affect downstream protein expressions, structures as well as their functions. It is suggested that different codon usage patterns should encrypt distinct characters for a certain type of organism, and as a result, a series of machine-learning models have been constructed, not only for learning the patterns from certain species, but also for predicting the species based on given patterns. Two gene segments of influenza A virus, hemagglutinin (HA; gene 4) and neuraminidase (NA; gene 6), were so essential for the immune response of their hosts, that the serotypes of the viruses are named after their combinations. They thus become the objects of this study, and those proposed models work quite well on the designated tasks.

## Introduction

Influenza A virus (IAV), a member within the family of *Orthomyxoviridae*^*1*^, was not only regarded as a common cause of respiratory infections in many different animals, but also one of the most important infectious diseases for us, since it caused several great influenza pandemic in human history. It has a negative-sense, single-stranded RNA genome divided into eight segments, two of which encode the surface glycoproteins - haemagglutinin (HA) and neuraminidase (NA). Both proteins are the key elements for standard genotypic classification because they are also targets for development of vaccines and antiviral drugs^2^.

A specific amino acid (except Met and Trp) could be encoded by a bunch of codons because of the redundancy, and those codons responsible for the same amino acid are called synonymous codons. Codon usage is neither uniform within nor between species despite the fact that synonymous codons don’t actually alter the amino acid sequences, thus species - specific codon usage bias^3,4^ would be observed instead of a uniform pattern. The choice of preferred codons over others within the same synonymous group reflects the distinct evolutionary forces that shapes the organisms and determines their overall fitness via a series of processes such as RNA processing, gene expression control as well as protein translation and folding steps^5–7^.

Many previous studies were trying to find out the reasons behind, and most of their conclusions indicated that mutation or natural selection could be two independent forces responsible in molecular evolution^5,8,9^. The explanations involving mutational mechanisms described a neutral fitness state owing to the underlying mutational properties, such as point mutations, contextual biases in the point mutation rates, and biases in repair^10^. Whereas explanations of nature selection thought in an opposite way that synonymous mutations could either promote or regress the fitness of an organism throughout evolution^11^. These two types of mechanisms were able to work solely or even together for shaping the synonymous variations among genomes and leaving some patterns that can be distinguished between species. For this reason, it is suggested that advanced mathematical models are needed to fit and learn those complicated features.

Machine learning is a very powerful tool for classification and error detection, and conventional machine learning algorithms, such as Support Vector Machines (SVM), Hidden Markov Models and Bayesian Networks, have already been widely adopted by bioinformatics^12^. Yet, their performances rely highly on the features that designed and extracted by human^13,14^. Deep learning, which belongs to the branch of supervised machine learning, has an architecture consists of multiple layers of nodes (or “Neurons”) for automated feature extraction and better scalability, since high level of abstraction can be learned thanks to the nonlinear transformation along the layers^15^. Convolutional Neural Networks (CNN) is a special type of Feed-forward Neural Networks which has been increasingly popular these days^16,17^. It was initially designed for image processing^18,19^ and was regarded as a simplified Neocognitron model that was proposed to simulate the human visual system^20^ with three main types of neural layers: convolutional layers, pooling layers, and fully connected layers. In addition to its aptitude of detecting local patterns such as edges and curved lines in the images, it is still doing better than Simple Feed-forward Neural Networks in some ways because it could be regarded as the latter model using a sparse weight matrix, which reduces the number of the network’s tunable parameters and thus increases its generalization ability^21^.

Here, we constructed a series of machine learning models on the HA and NA genes of several representative influenza A virus serotypes, from the simplest Multinomial Logistic Regression, to the Simple Feed-forward Neural Networks, and finally a more complicated Convolutional Neural Network (CNN), in order to capture the codon usage features beneath different serotypes of influenza A viruses. By applying these models, the distinct evolutionary trajectories of influenza serotypes are supposed to be better reflected and predicted.

## Results

### 1) Principle component analysis (PCA) for analyzing the degree of data concentration

As we can see from Figure 1.1, the first and second principle component (PC1 and PC2) of the original 64-dimension RSCU values can explain close to 50% of the total variation (PC1 explained 24.61% whereas PC2 explained 19.44%, and both add up to 44.05%). However, the impact of the third and latter components don’t drop sharply, giving a long tail in the plot. This means the variation of the third and latter component also have their own importance. In order illustrate this issue in greater details, we plot PC1 against PC2 in the same plot in Figure 1.2. Here we can see that using these two major components for visualization, the samples of 15 serotypes did aggregate in groups, but instead of having very clear boundaries, some of the dots in this dimensional reduction plot do stack against each other. According to the purposes of separating those 15 serotypes, models that can utilize all of the 64 codon usage dimensions would be constructed.

**Figure 1.1.**
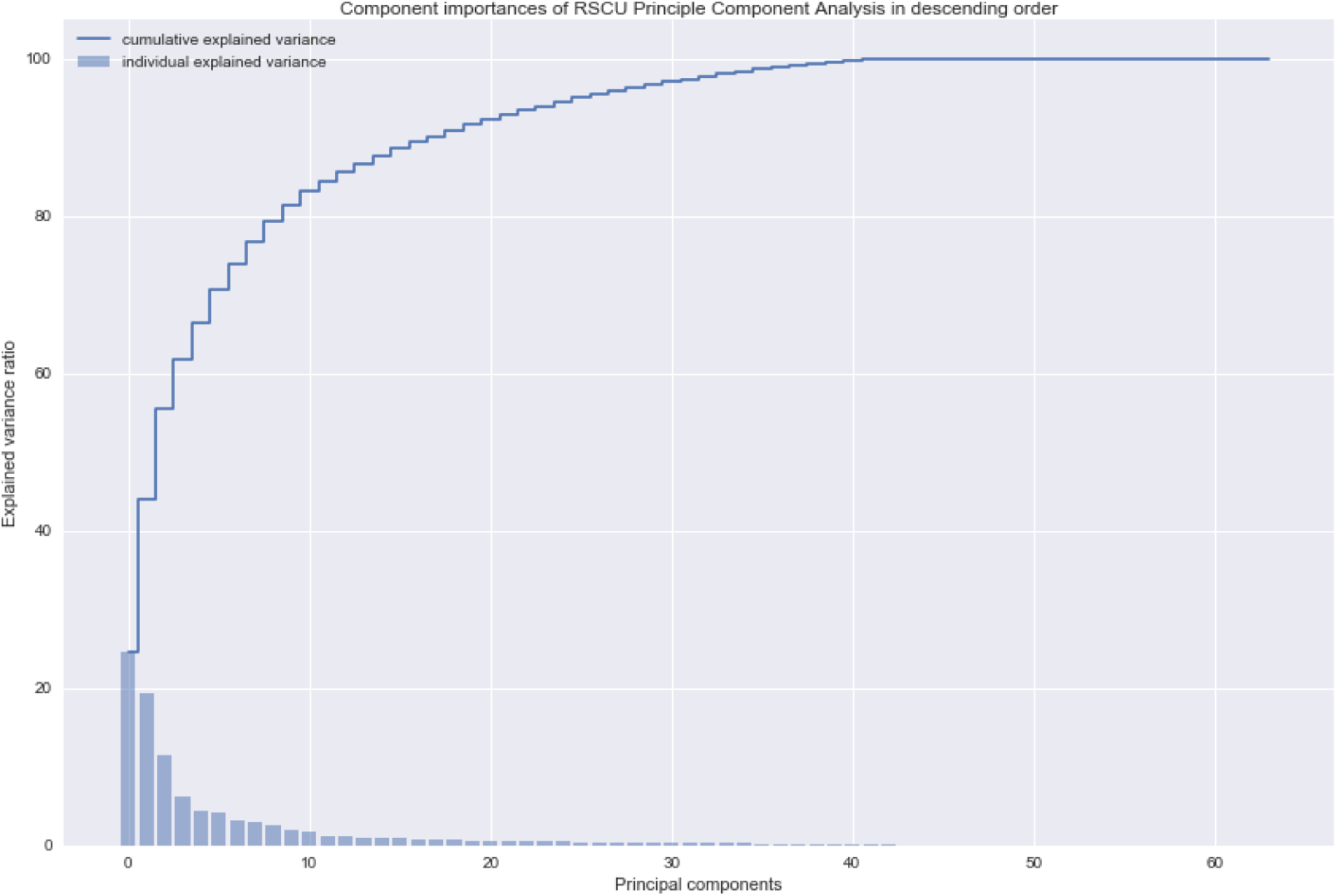
The contribution of every principal component to the total variance after PCA (Principle Component Analysis) on 64 RSCUs. Bar plot represents the results of them each, and the line chart represents their cumulated results.

**Figure 1.2.**
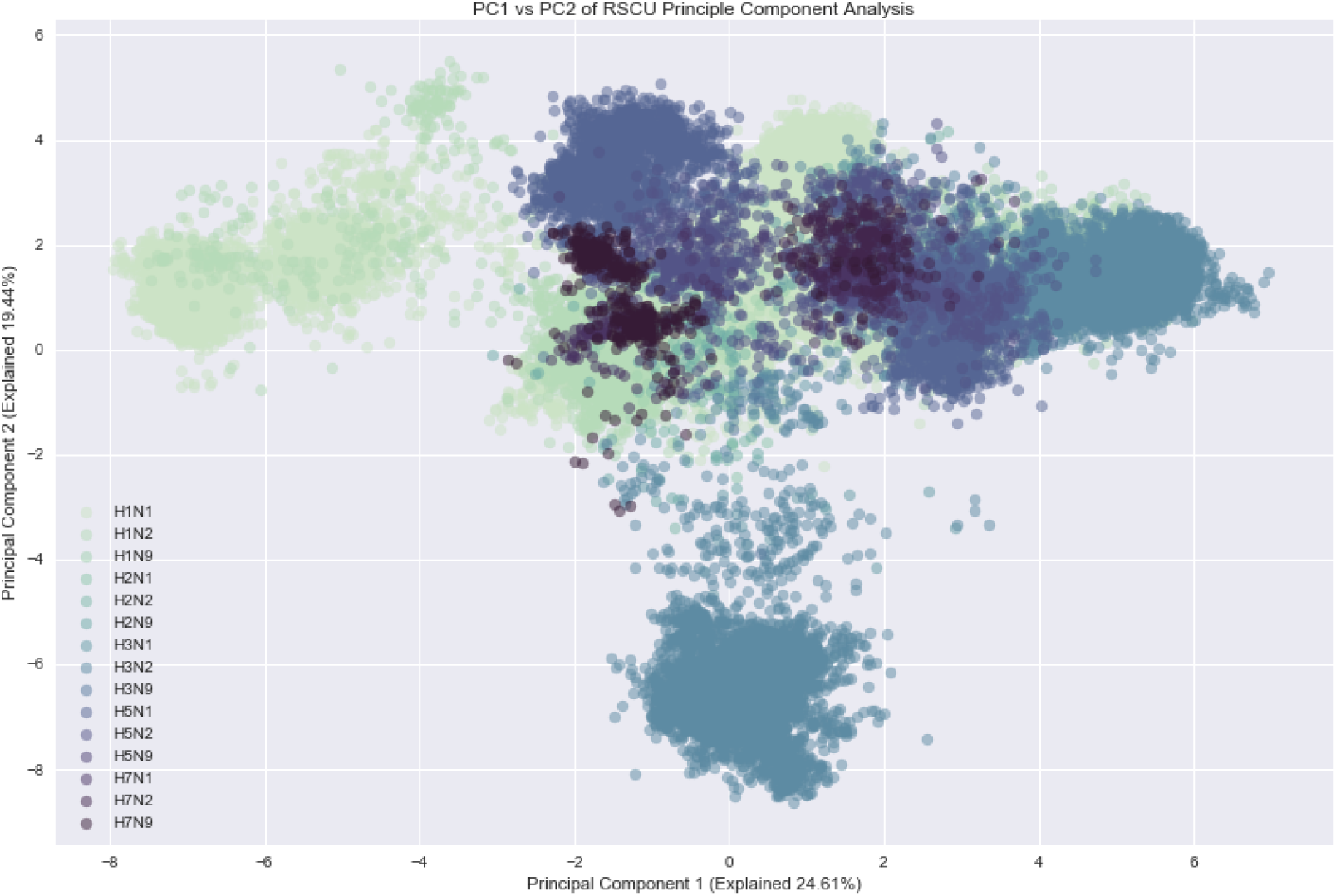
PC1 vs PC2 after PCA (Principle Component Analysis) on 64 RSCUs. Every single dot represents the result of a sample (either from HA or NA gene), and different serotypes are represented by different colors.

### 2) A simple Multinomial Logistic Regression for starting point of the model

A good starting point is the Multinomial Logistic Regression model (or Softmax Regression in some of the machine-learning literatures, a typical model for multi-class classification problems), not only because of its simplicity (Figure 3.1 clearly demonstrates its design), but also because it has a comprehensive parameter structure, so that people could visualize and understand it with ease.

Figure 2 shows the distributions of its final parameters (the model achieves an overall accuracy score of 87.9% on the test dataset when 4000 iterations of training has been done) and from this plot, we discover that different serotypes will lay particular emphasis on different codons. For instance, H1N1 regards UUA as an important codon because it is given a very high weight (the color of the corresponding box bias to crimson), and H3N1 regards UCA as very unimportant for its low weight (the color of the corresponding box bias to Mandarin blue). Despite its simplicity and relatively low accuracy, this model did successfully extract some pattern from 64 RSCU combinations of those 15 serotypes. To illustrate the powerfulness of using such a model, each codon can plot 15 different RSCU distributions according to their serotypes, yet the individual patterns or their combinations can hardly be distinguished by human’s eye (Figure S1).

**Figure 2.**
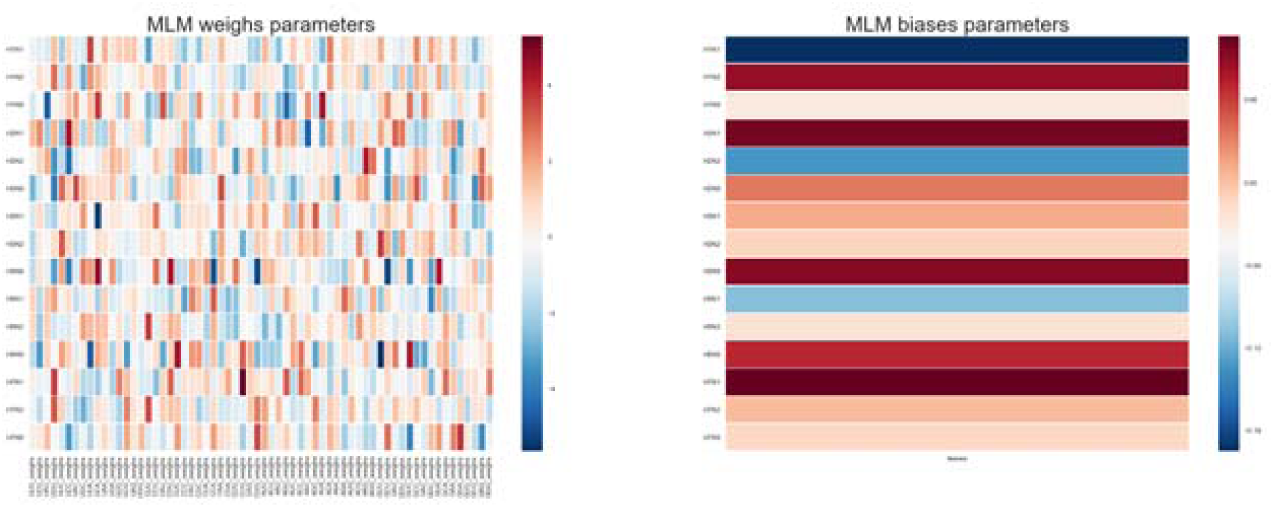
The parameter distribution of Multinomial Logistic Regression model. Left: weighting parameters; Right: bias parameters.

The result here also explains the reason of using all of the 64 codons: traditionally, the study of codon usage biases will use either 61 codons (exclude 3 stop codons) or 59 codons (further exclude the UGG and AUG, because these two has one-to-one corresponding relationships with the amino acid they have encoded). The parameters for three stop codons (UAA, UGA, UAG) and two other codons (UGG, AUG) are very close to zero (the colors of the corresponding boxes bias to a light color), giving a sign of their insignificance for the model. For this reason, we can keep all those 64 dimensions for latter models (2D and 3D Convolutional Neural Networks) which require orderly input shapes of 4×4×4.

### 3) Complex models adding nonlinearity

To investigate if there is any room for improvement, the models below are constructed via a series of thinking:

First, based on Multinomial Logistic Regression, more matrices are added and they are connected by activation functions. This gives the Feed-forward Neural Networks, which would have more parameter spaces and more importantly, more nonlinearity.

Second, inspired by image recognition tasks (basic elements such as lines and dots are the building blocks of a picture, thus the values of nearby pixels would affect the performance of a model together other than separately), 2D and 3D Convolutional Neural Networks are modified and applied to study the codon usage pattern. Because image data often come with a rectangular (image with only gray level) or cuboid shape (image with RGB color channels), each set of the 64 RSCU values are converted to a 4×4×4 cube before feeding into the models. These two models not only have the nonlinearity as the Feed-forward Neural Networks, but also can detect the patterns between adjacent elements. Figure S3 shows the configuration of the 4×4×4 cube. For the depth axis, nearby codons will have a 1-bp difference in the 1^st^ position (e.g. AAA-CAA-GAA-UAA). For the height axis, nearby codons will have a 1-bp difference in the 2^nd^ position (e.g. AAA-ACA-AGA-AUA). For the depth axis, nearby codons will have a 1-bp difference in the 3^rd^ position (e.g. AAA-AAC-AAG-AAU). Reshaping data like this could represent the effects of 1-bp mutational change of among codons, and in this design, 2D Convolutional Neural Networks can detect mutational pattern of 2^nd^ and 3^rd^ position, whereas 3D Convolutional Neural Networks can detect all of the three.

(1) Feed-forward Neural Networks: give more non-linearity for Multinomial Logistic Regression Feed-forward Neural Networks, which can be regarded as many layers of Multinomial Logistic Regressions stacked together (Figure 3.2 shows the structure) with the non-linearized ReLU functions in between, does give out a better overall accuracy score of 96.7% on the test dataset. It could be learned from those above that, in order to separate different codon usage patterns with 64 dimensions, a classification model with the capability of twisting the hyper planes between scattered sample values in hyperspace seems to give a much better result, than the one only using linearized hyper planes, even though the parameter distribution of this new model is not easy for people to interpret intuitively (Figure S2.1-S2.3). Sometimes a model with a higher accuracy score is not equivalent to a better overall performance, thus we apply some other indexes such as the overall recall (72.5 % vs 70.7%), precision (68.5% vs 50.9%), F1-score (70.0% vs 54.6%) and MCC coefficient (95.2% vs 83.3%), all of which are better for the Feed-forward Neural Networks than in Multinomial Logistic Regression. As a result, the improvement here is rock solid since no compromise is observed among accuracy, recall and precision on the more complex Feed-forward Neural Networks.

**Figure 3.1.**
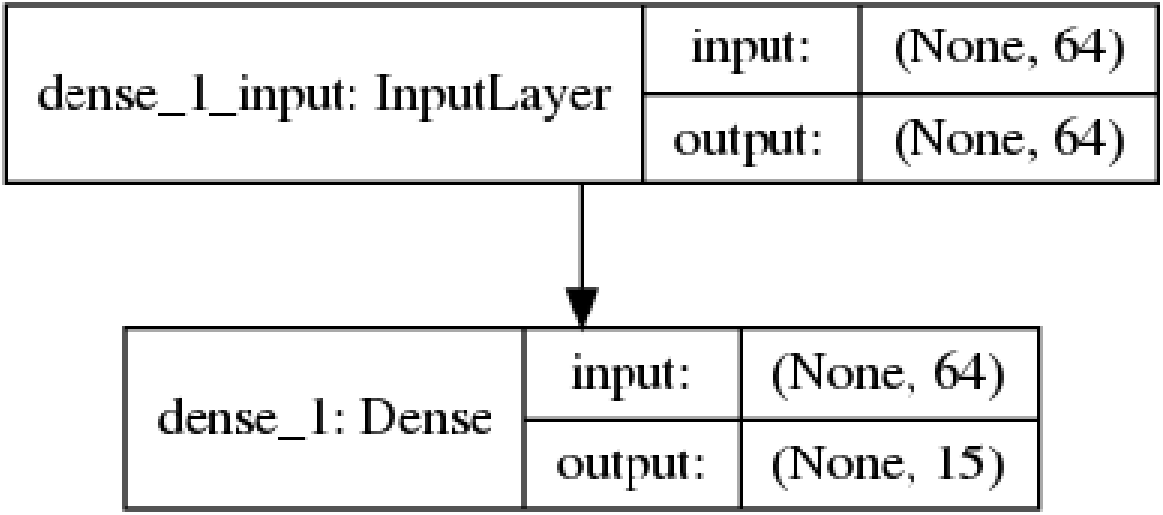
The structure of Multinomial Logistic Regression model.

**Figure 3.2.**
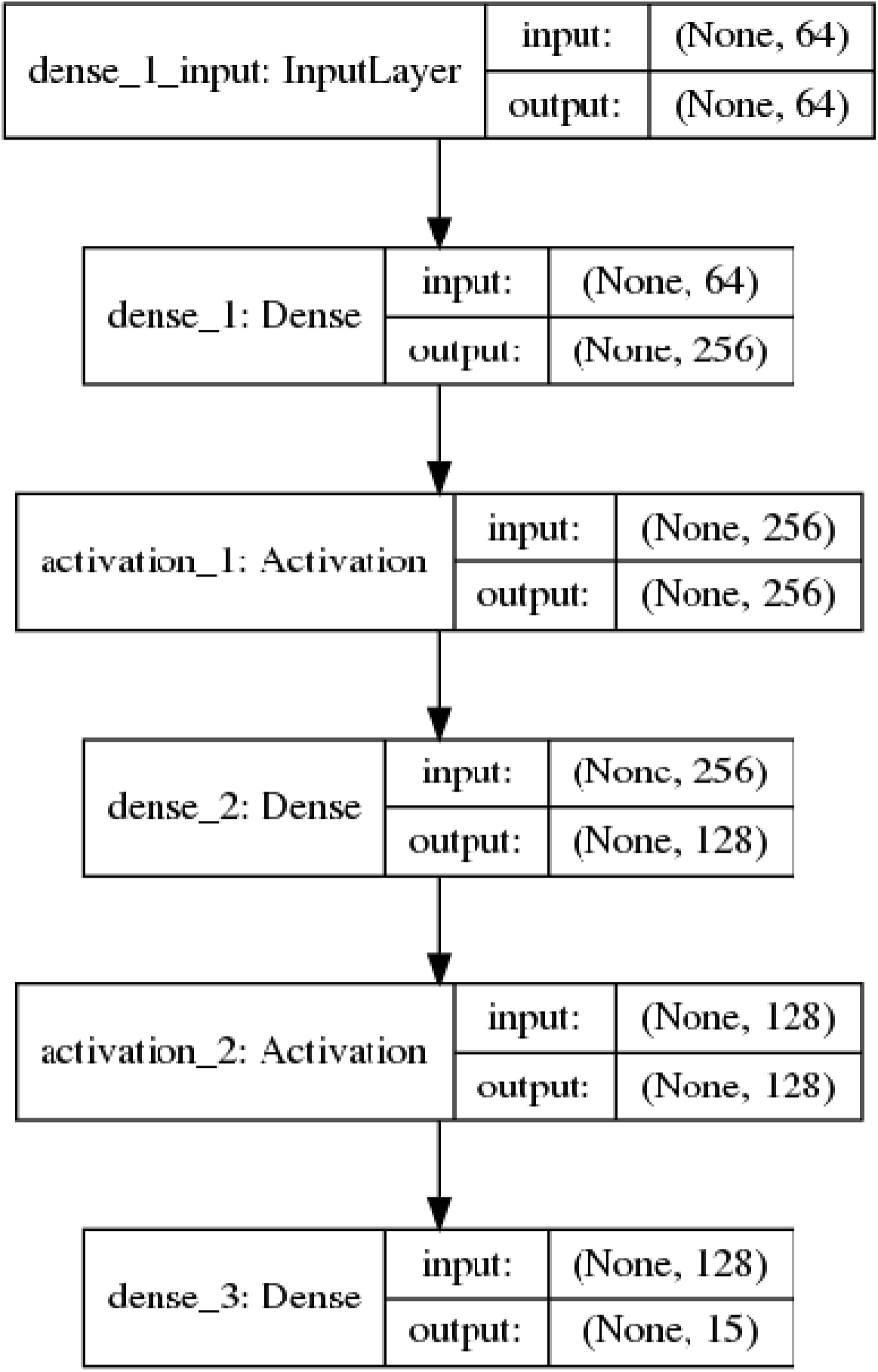
The structure of Feed-forward Neural Networks model.

**Figure 3.3.**
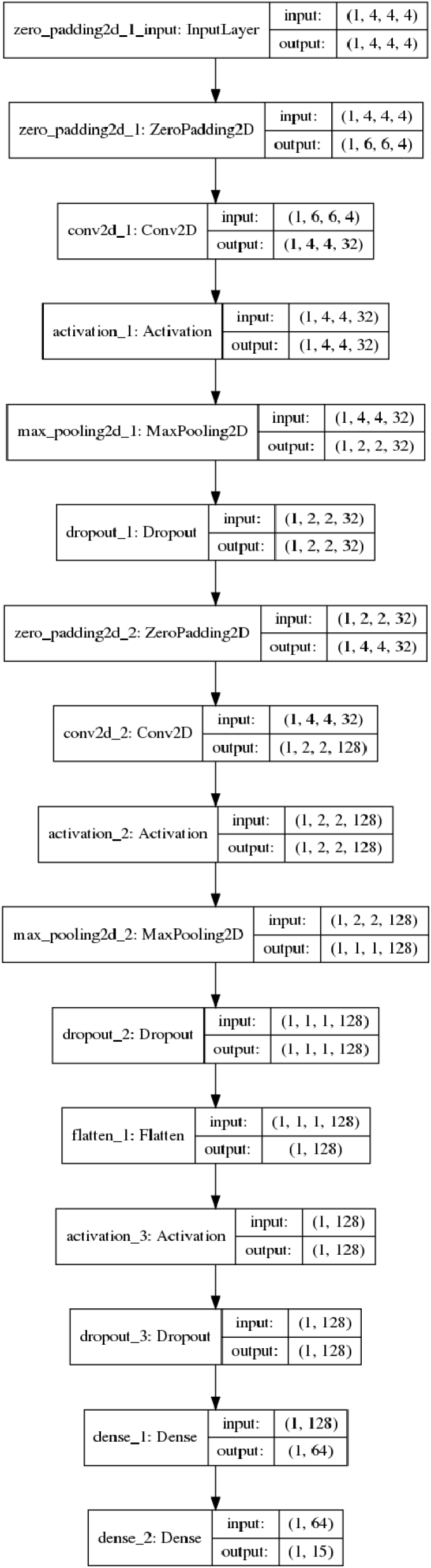
The structure of 2D Convolutional Neural Networks model.

**Figure 3.4.**
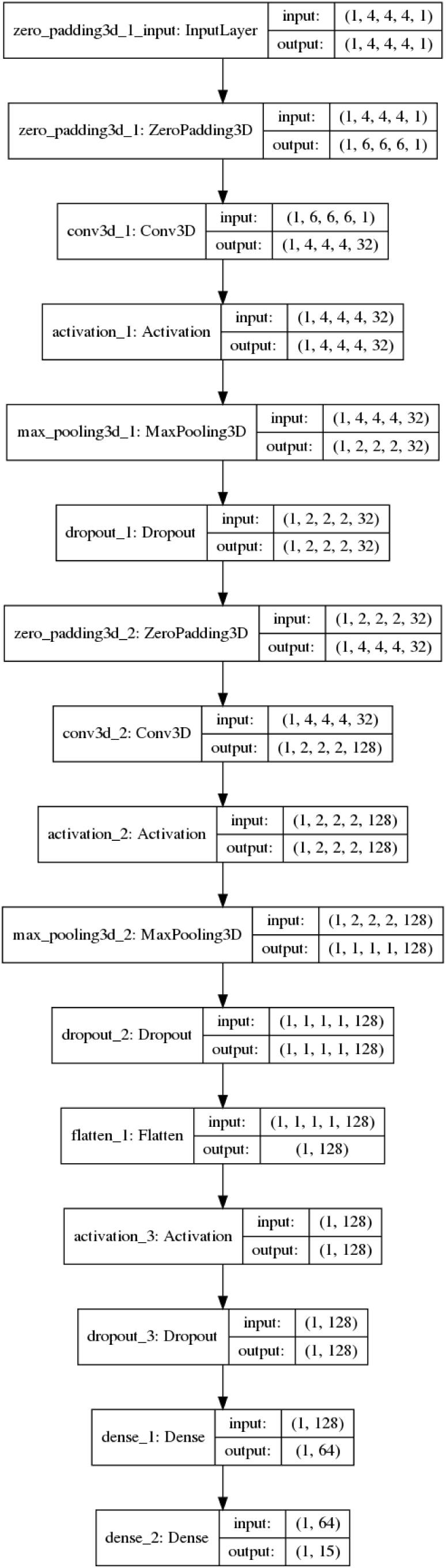
The structure of 3D Convolutional Neural Networks model.

(2) 2D and 3D Convolutional Neural Networks: detecting neighboring effects fora specified 4×4×4 data structure

Both 2D and 3D Convolutional Neural Networks have been done (see Materials and Methods for details, and Figure 3.3 and Figure 3.4 for their structures), and accuracy scores of 94.8% and 97.0% had been achieved. Since these two models have more assumptions than the Feed-forward Neural Networks, a thorough comparison will be made in the next part.

### 4) Pros and Cons for different models

An overall performance comparison is made between the models (Figure 4.1, exact values refer to the table of Overall Benchmarks), which shows that the model of 3D Convolutional Neural Networks ranks the top, and Multinomial Logistic Regression ranks the bottom. Feed-forward Neural Networks and 2D Convolutional Neural Networks lie between these two, but the former one seems to perform better.

**Figure 4.1.**
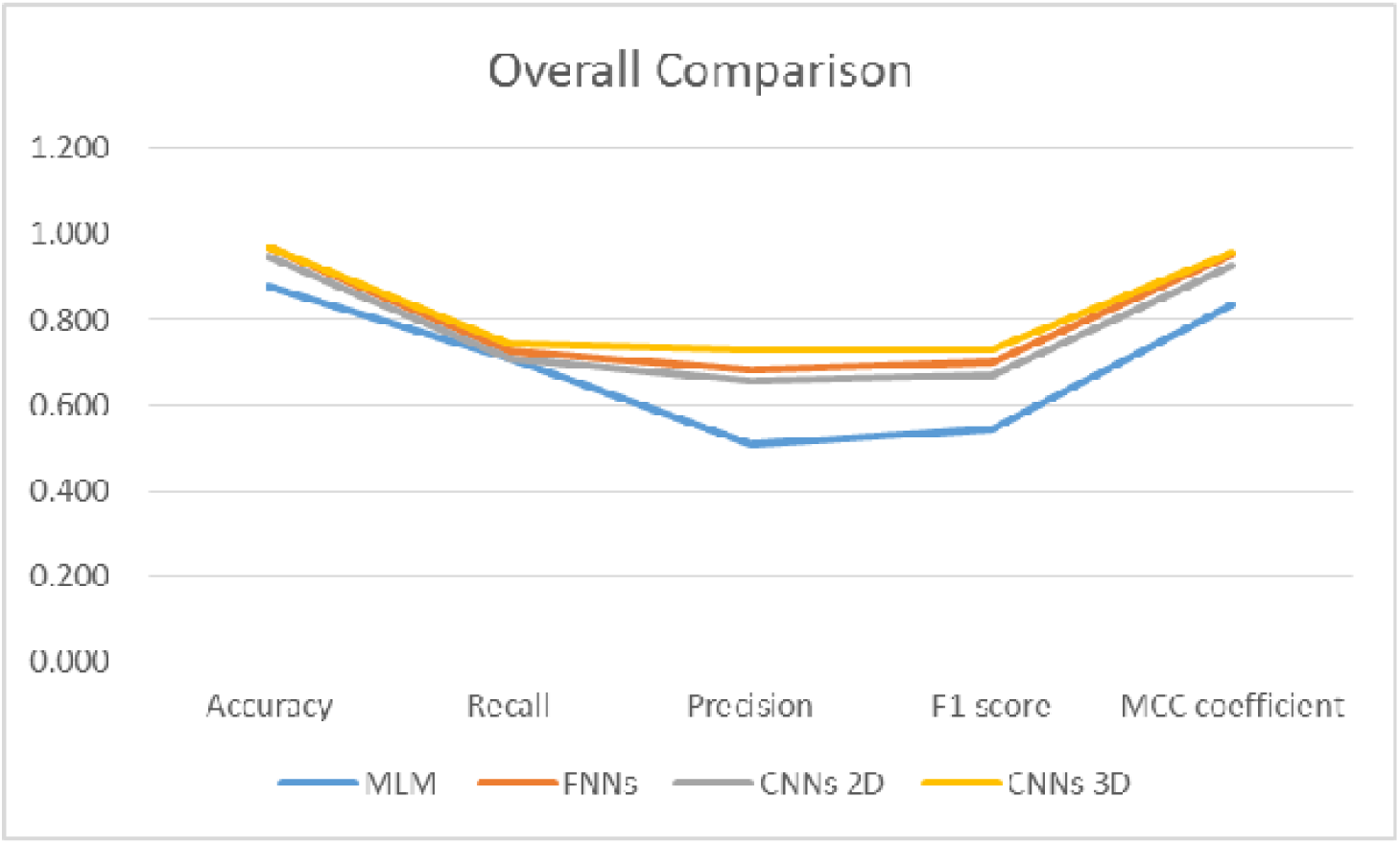
Parallel comparison for the evaluation indexes of different models on test set. Those numbers are based on the collections of all serotypes.

A possibility is proposed: the difference of 2D and 3D Convolutional Neural Networks, according to our construction, is that it the 2D model can detect codon difference patterns in 2^nd^ and 3^rd^ position, whereas 3D model can detect all of the three: 1^st^, 2^nd^ and 3^rd^. As the matrix constructed here for RSCU values represents the similarities between the codons, this means the effect among adjacent elements should not be ignored. A change at the 1^st^ and the 2^nd^ position of a codon will always be a non-synonymous one for the amino acid, whereas the 3^rd^ position will mostly be a synonymous one. If there is no selective constrains over genes, codons will be freely mutated and the matrix will have same value across 64 positions. If external forces are exerted on genes, the ratios between each of the 64 positions will be changed, and the final result of the matrix can be regarded as distinct features for each gene. Missing the ability of detecting the patterns in the 1^st^ position, an essential part of the codon usage patterns, could result in the performance difference between 2D and 3D model. 2D model also performs worse than the Feed-forward Neural Networks, because mathematically, it has less connections between its neurons (Computational result of each neuron in one layer of Feed-forward Neural Networks is affected by all of the neurons in the last layer, and will affect all of the neurons in the next layer. Each neurons of Convolutional Neural Networks, however, will affect or will be affected by nearby neurons in the next or last layers).

It should be noticed that 3D Convolutional Neural Networks has fewer parameters compare with the Feed-forward Neural Networks (Multinomial Logistic Regression: 64+15=79; Feed-forward Neural Networks: 64×256+256+256×128+128+128×15+15=19087; 2D Convolutional Neural Networks: 3×3×32+32+3×3×128+128+128×64+64+64×15+15=10831; 3D Convolutional Neural Networks: 3×3×3×32+32+3×3×3×128+128+128×64+64+64×15+15=13711), even though its performance is superior to the latter. Yet, trade-offs still exist, for this model takes so much time to train (Multinomial Logistic Regression and Feed-forward Neural Networks take less than one day, 2D Convolutional Neural Networks takes about two days, and 3D Convolutional Neural Networks takes more than a week, in the same computer platform).

Detailed comparisons among serotypes give more thorough analysis on this standpoint: the overall trend is similar as it is above, but there are still a few exceptions (Figure 4.2-Figure 4.5). Serotypes with large sample numbers do support this very well (e.g. in the test set, the top four serotypes are H1N1: 5773, H1N2: 1035, H3N2: 5733, H5N1: 1170). Multinomial Logistic Regression can sometimes do very well (e.g. H5N1 have both very high accuracy and F1 score), but other times it tends to fall way behind those complex models because it has to make a compromise between accuracy, precision and recall values. An investigation is made on the sub table of “serotype based compare” in the supplementary excel file “Benchmarks”. For each model, accuracy is plot against F1 score and each dot represents the performance of a certain serotype. In complex models, the correlation between those two using R^2^ is very obvious, but in Multinomial Logistic Regression, it tends to be very low. It seems that the complex models could raise both the accuracy and F1 score in the same time. By contrast, if simple model wants to lay particular emphasis on one side, it has to make a concession on the other. Serotypes with medium sample numbers continue this trend (e.g. in the test set, four serotypes with medium sample numbers are H2N2: 89, H5N2: 303, H7N1: 81, H7N9: 143), but the performances may not be as good as the above. For serotypes with small sample numbers, however, simpler model could achieve much better accuracy values than complex models, despite the fact they have to compromise a lot in other values such as precision, recall and F1 score (e.g. H1N9, H3N1, H5N9). This is owing to the data bias and it will be discussed in the later section. Yet, H7N2 is an exception, since the performances of those models on this serotype are rather good.

**Figure 4.2.**
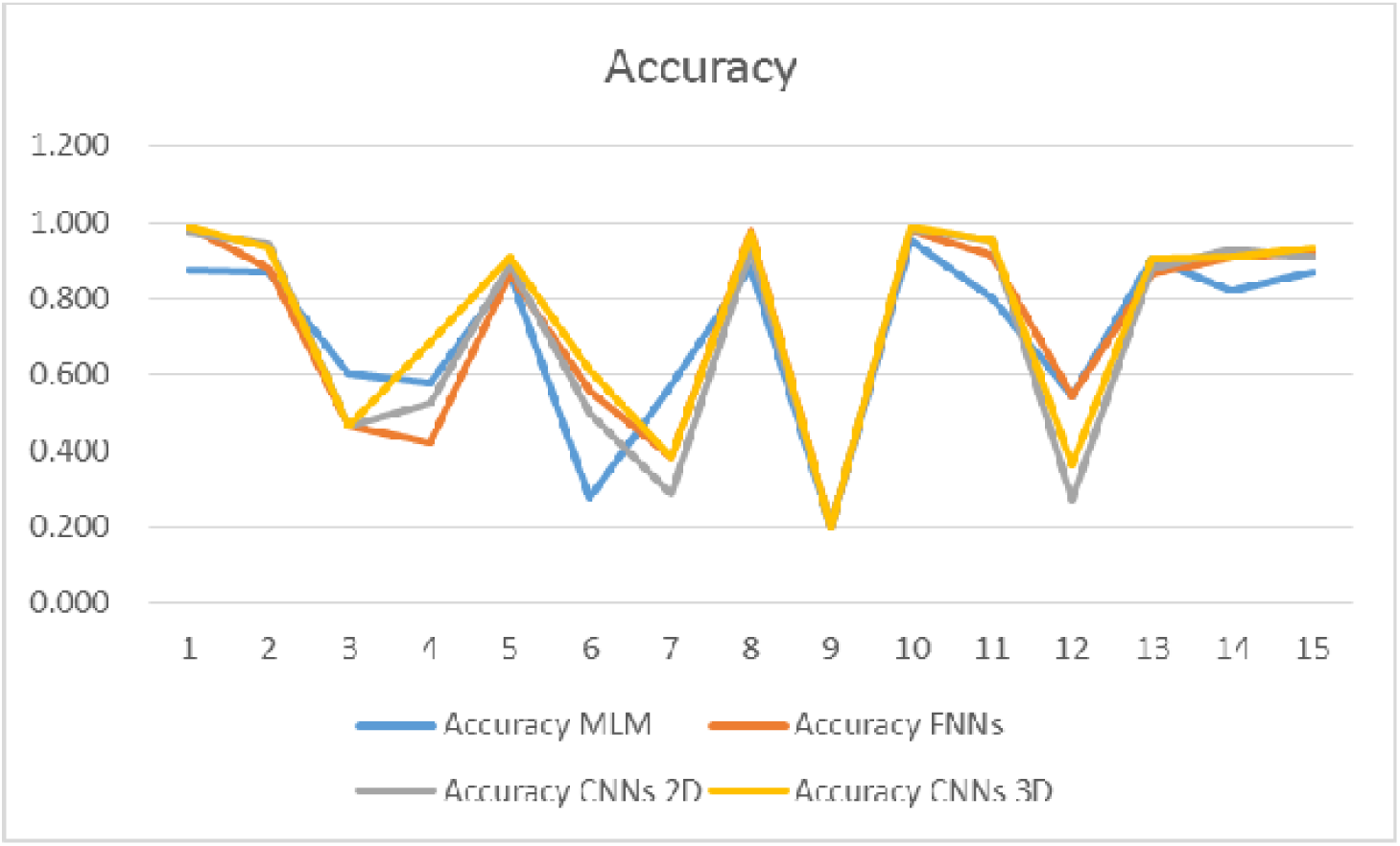
Parallel comparison for the Accuracy of different models on different test set based on serotypes. Number 1-15 denotes: H1N1, H1N2, H1N9, H2N1, H2N2, H2N9, H3N1, H3N2, H3N9, H5N1, H5N2, H5N9, H7N1, H7N2, H7N9.

**Figure 4.3.**
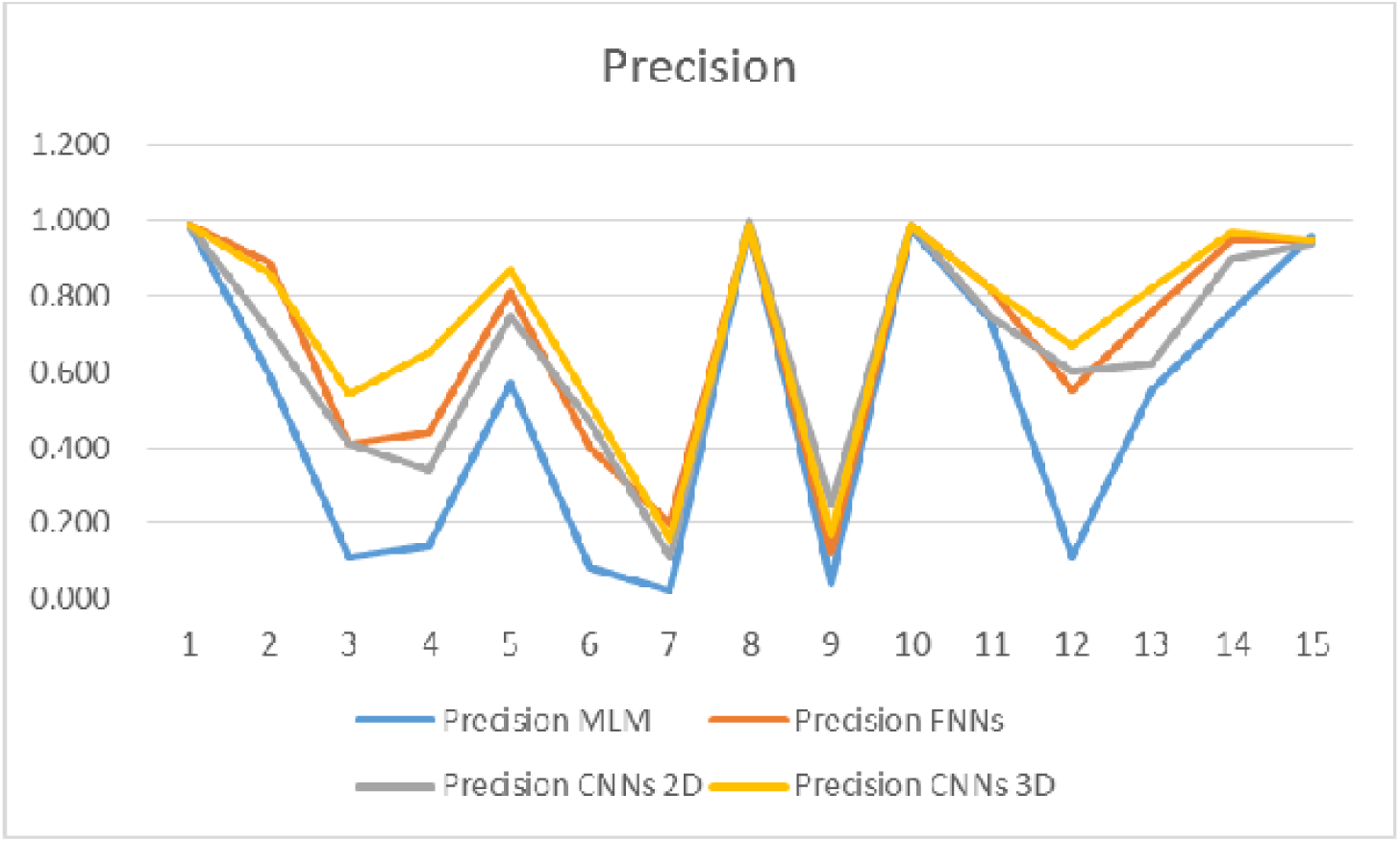
Parallel comparison for the Precision of different models on different test set based on serotypes. Number 1-15 denotes: H1N1, H1N2, H1N9, H2N1, H2N2, H2N9, H3N1, H3N2, H3N9, H5N1, H5N2, H5N9, H7N1, H7N2, H7N9.

**Figure 4.4.**
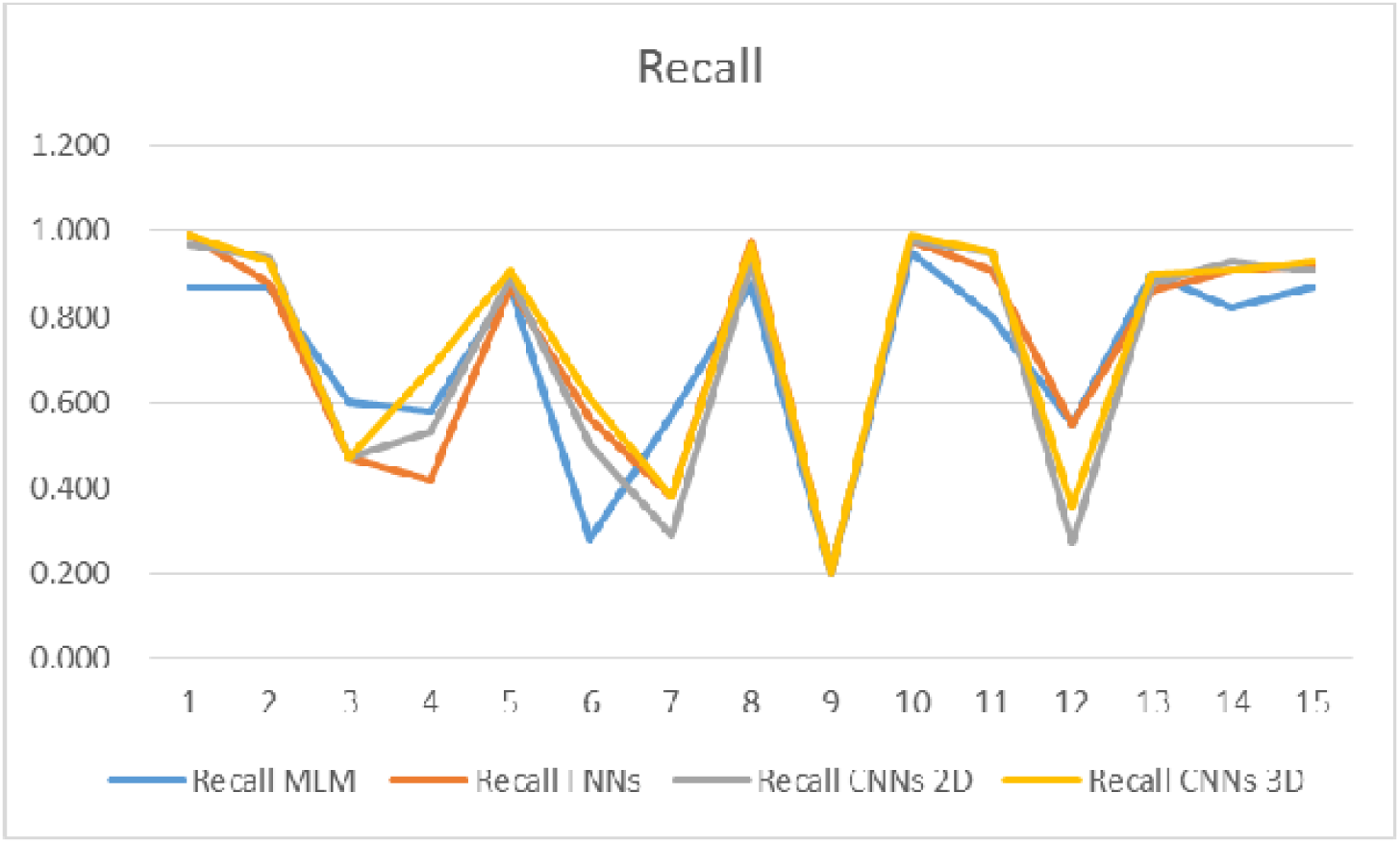
Parallel comparison for the Recall of different models on different test set based on serotypes. Number 1-15 denotes: H1N1, H1N2, H1N9, H2N1, H2N2, H2N9, H3N1, H3N2, H3N9, H5N1, H5N2, H5N9, H7N1, H7N2, H7N9.

**Figure 4.5.**
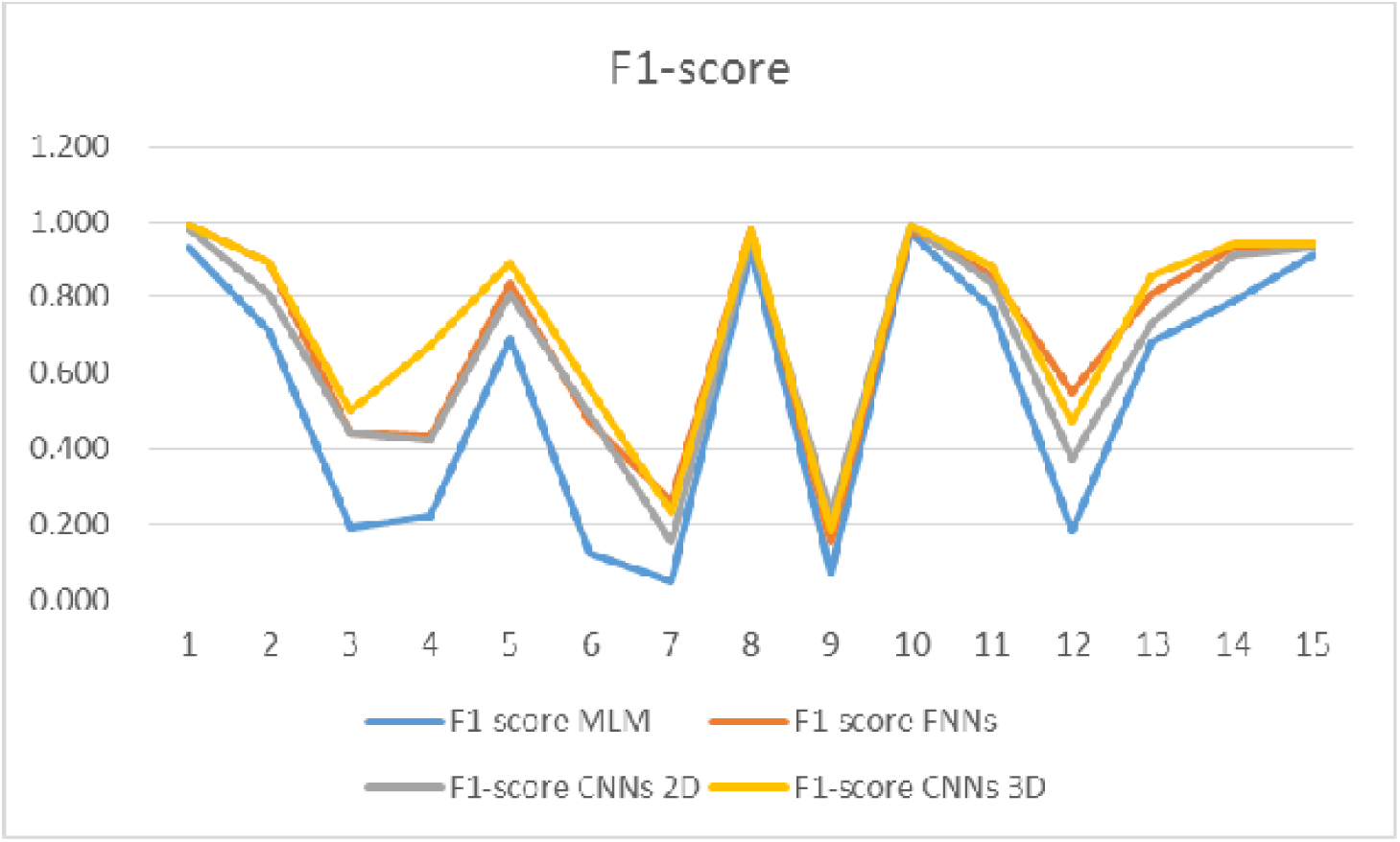
Parallel comparison for the F1 score of different models on different test set based on serotypes. Number 1-15 denotes: H1N1, H1N2, H1N9, H2N1, H2N2, H2N9, H3N1, H3N2, H3N9, H5N1, H5N2, H5N9, H7N1, H7N2, H7N9.

**Figure 4.6.**
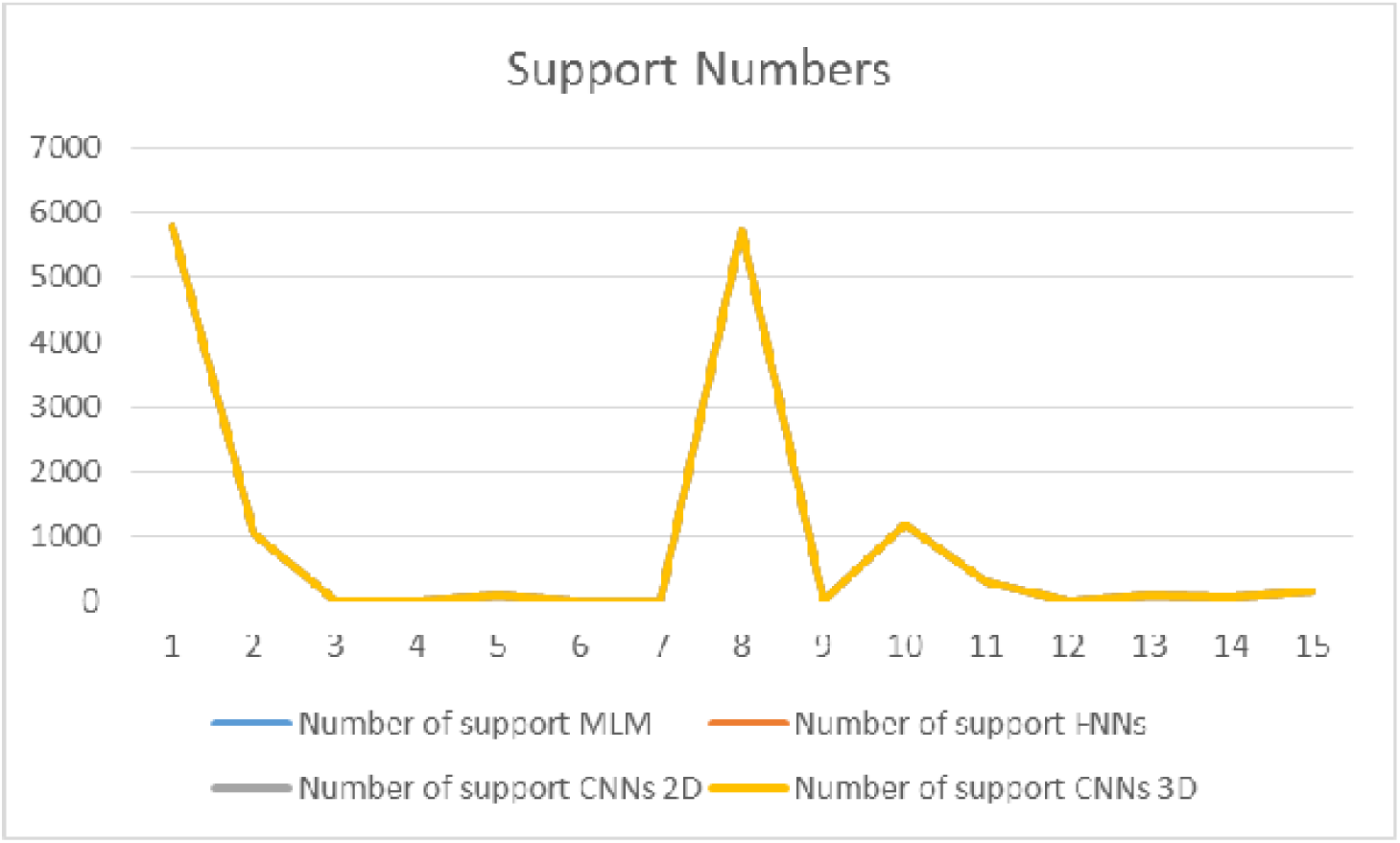
Support number difference for test set based on serotypes. Number 1-15 denotes: H1N1, H1N2, H1N9, H2N1, H2N2, H2N9, H3N1, H3N2, H3N9, H5N1, H5N2, H5N9, H7N1, H7N2, H7N9.

### 5) Excluding the artifacts of the models: negative results (continents as tags) and small sample numbers

Some artifacts should be ruled out when using these models. Here are some additional analysis for discussing these problems.

The first one is to prove that these models did get biologically meaningful information according to the assumptions have been made in advance, and that is: they could accurately reflect the congruent relationship between codon usage pattern and its corresponding serotype. To prove this, instead of using “serotypes” as labels, negative samples have been created using the same genes with the same RSCUs values as input but “continents” as the alternative tags (The influenza database has samples from these six continents: Africa, Asia, Europe, North America, Oceania, and South America). After training these models for the same 4000 iterations with very similar constructions (only output layers have been modified to adapt to the number of output), on the separate test set (still adopt the 7:3 train-test split), the Multinomial Logistic Regression, Feed-forward Neural Networks, 2D and 3D Convolutional Neural Networks achieve accuracy scores of 33.3%, 69.5%, 60.5%, 70.3% (See the table: Benchmarks – continents, for details). Contrary to the models trained using serotype tags, these results are much worse than that. Thus, we believe that codon usage patterns should be existed in different serotypes.

The second one is owing to the offset of the samples with different number of tags or labels. Using the test set as an example (the training set will have a similar proportion), H1N1 has the largest support number of 5773 while the H3N9 has the smallest of 5. Their F1-scores also have huge differences (Multinomial Logistic Regression: 93% vs 7%; Feed-forward Neural Networks: 99% vs 15%; 2D Convolutional Neural Networks: 98% vs 22%; 3D Convolutional Neural Networks: 99% vs 18%), despite the fact that data augment method (SMOTE) have been done. It is suggested that the eagerness for a large amount of data, a universal character for machine-learning models especially those complex ones, might be responsible for this phenomenon. To prove this, we plot the support number against accuracy or F1 score on each model, and it is very clear that they do have some correlation (See the sub table of “serotype based compare” in the supplementary excel file “Benchmarks”). A separate training and testing process is then added only using those serotypes with big numbers. As expected, much better overall performances have been achieved (See the supplementary excel file “Benchmarks - only large samples” for details).

## Discussion

Primarily, the benefit of building up a machine-learning model, in contrast to the unsupervised approaches such as clustering and dimensionality reduction widely adopted in previous studies of codon usage biases, is that it can do several things altogether: to learn and to predict. In other words, not only can it summarize codon usage patterns and group together the similar ones based on current data, but also being able to generalize those learned information and infer new unknown samples via these models as well. Technically, the codon usage RSCUs of the influenza A viruses here have been randomly shuffled and split into training set and test set with a ratio of 7:3. Indeed, they are two independent parts coming from the same distribution, so that the test set would correctly justify the performances of the models. Thus, if the codon usage patterns of a virus strain don’t change too much over time, we can still do a proper prediction based on the historic patterns.

Secondly, when training these models, rather than joining HA and NA genes together from the same virus sample and form a 64-dimensional RSCU vector, either one of them is fed to the model one at a time albeit the whole training set is a mix of these two. As a matter of fact, these models are quite conclusive because they should make a balance between these two defining components for correctly predicting the influenza serotypes labels, and they still deliver reasonable outcomes in the test set. And indeed, a clear pattern is given since a very strong connection between the codon usage patterns of HA or NA gene and its corresponding serotype has been built.

Mathematically, this is very close to an “end to end” machine-learning for codon usage study, because these models take information directly from all 64 RSCU values instead of only a few of the important ones, and reducing the possibility of introducing error or losing information when using summarized data.

Biologically, codon usage pattern here is regarded as a kind of innate feature belongs to a certain type of organism (just like fingerprints or iris to human). As we demonstrate in the result section that, according to our purposes of distinguishing different serotypes, their features cannot be represented well by simply visualizing the distributions of their 64 individual components, or using a conclusive approach of dimensional reduction like Principle Component Analysis (PCA). In contrast, those machine-learning models do give satisfactory results.

In addition, an innovation is made during the pre-processing procedures of 2D and 3D Convolutional Neural Networks, because it includes the reshape of 1D codon usage vector to a 4×4×4 3D matrix with highly correlated nearby elements. Since the application of Convolutional Neural Networks inspired from image recognition tasks also gives a satisfactory result here, it provides new ideas for how to manipulate biological data, so that their data structures will be able to adapt to some very advanced and effective machine learning models.

Based on the work of influenza A virus, we might expand it to the field of cancer biology in the near future, where there are much more complicated patterns (chromosome patterns, sequence patterns, mutation patterns, etc.) for us to conquer. Also, back to the codon usage problems themselves, we might try to combine more genes (denote as number “n”) and their 64 RSCUs (denote as number “64”) in the next study, so that the input could be in the shape of n*64. Models like Convolutional Neural Networks would be further modified and tested in order to see if there is any pattern.

In a more general sense, this study sets up a new starting point of analyzing genome data.

## Materials and Methods

### Material

Gene sequences of hemagglutinin (HA) and neuraminidase (NA) of influenza A viruses (IAVs) used in this study are downloaded from NCBI Influenza Virus Resource (https://www.ncbi.nlm.nih.gov/genomes/FLU)^22^. Here are the reasons of choosing these subtypes. Even though the number of possible IAVs combinations is huge, given the fact that there are 18 types of HA genes and 11 types of NA genes (till the year of late 2017, from NCBI), it is interesting that very few have been reported to infect humans^23^. Common subtypes, such as H1N1, H1N2, H2N2, H3N2, and H5N1^24–27^, were found to circulate around humans to cause seasonal flu. In addition, some of them even cause great pandemics in the human history (e.g. H1N1 in 1918, H2N2 in 1957, H3N2 in 1968)^28–29^. Recent reports show that the avian origin H7N9 might have a pandemic potential^30^, so it has been considered as well. Thus, 5 HA types (No.1, 2, 3, 5, 7) and 3 NA types (No.1, 2, 9) have been chosen with all their combinations (15 combinations in total). Also, all of their possible hosts have been included to see if the model are generalized.

### Method

#### 1) Data Normalization: Relative synonymous codon usage (RSCU) analysis

Numbers that are fed to those models should be transferred to a similar scale, and to be specific, the raw numbers of codon usage must be standardized against the length of gene sequence and the frequency of the amino acid. Sharp & Li designed a relative synonymous codon usage (RSCU) index assuming a reference state that all codons for a particular amino acid are used equally, and then these expected numbers are compared with the actual observed numbers in the form of the following equation^31^:

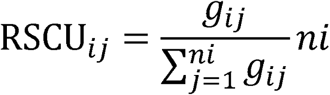

g_ij_: Observed number of the j^th^ codon for the i^th^ amino acid

ni: Number of synonymous codons that the i^th^ amino acid has.

Interpretation of RSCU value:

RSCU value < 1.0: observed number less than expected, negative codon usage bias;

RSCU value = 1.0: no bias;

RSCU value > 1.0: observed number more than expected, positive codon usage bias.

#### 2) Model construction

Supervised machine learning algorithms are designed to build a classifier, which is capable of distinguishing several specified classes from the input vectors, by learning a set of training samples. Since different models require different shapes of input data, those 64 RSCU values of a single gene have been reshaped to a 1-D vector (1×64, for PCA, Multinomial Logistic Regression and Feed-forward Neural Networks) or a 3-D vector (4×4×4, for those two Convolutional Neural Networks).

The whole dataset has been randomly shuffled, and then was divided into training data and testing data with a ratio of 7:3. Since this dataset is imbalanced, for some of the classification categories might have much more objects than the other ones, the training data have been augmented using Synthetic Minority Oversampling Technique (SMOTE)^32^, which make different classes equal in number to avoid model training biases.

##### (1) Principal component analysis (PCA)

Clustering methods such as correspondence analysis (CA)^33^, principal components analysis (PCA), k-means analysis and other statistical techniques^34,35^ were widely used in many previous studies to classify and categorize codon usage variations. Here we used the statistical method of PCA^36^, which was previously used for summarizing the most important codon usage pattern from data^37^, as a prior visual representation before we constructed the following machine learning models. The input vector used here were 64 RSCU values (in fact 59 of them are effective and commonly used in previous studies, but here in this study, two codons that encode Methionine and tryptophan with one-to-one relationships, plus three stop codons, were just left as they were for fulfilling the 4×4×4 matrix required by latter models, and the reason that they could be used would be explained later in the result section) from each single gene (either HA genes or NA genes, regardless of their serotypes). This would map the multi-dimensional data to a lower-dimensional hyperspace for maximizing the variation along each new axis, in order to see the degree of concentration for the multi-dimensional data.

##### (2) Multinomial Logistic Regression

A Multinomial Logistic Regression model^38,39^ (MLM for short, or Softmax Regression in some of the machine-learning literatures) is going to train an m-class classifier (in this case m = 15, representing 15 serotype combinations) with n training samples, and the number of features that each sample has is d (in this case d = 64, representing 64 RSCU values), written in the form of a 1-D vector ***x*** *=* [x_1_, x_2_… x_d_] ^T^ ∈ **R**^**d**^. The class label is represented by a “1-of-m” (or “One-Hot encoding” in some of the machine learning literatures) encoding vector ***y*** *=* [y^(1)^, y^(2)^… y^(m)^]^T^. If a sample ***x*** belongs to the class i, then y^(l)^ = 1, otherwise y ^(l)^ = 0. Thus, n training samples could be represented as the data set ***D*** = {(***x***_**1**_, ***y***_**1**_), (***x***_***2***_, ***y***_***2***_) … (***X***_***n***_, ***y***_***n***_)}. The probability that **x** belongs to class i is given in the following function (binary problems (m = 2) is known as a Logistic Regression Model):

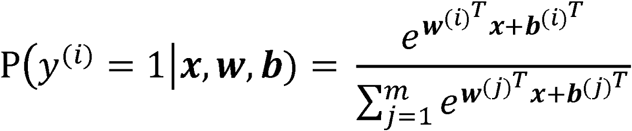

***w***^*(i)*^ weight vector corresponding to class i, for i ∈ {1, 2…m}, which has a dimension of d x m.

***b***^*(i)*^ : bias vector corresponding to class i, for i ∈ {1, 2…m}, which has a dimension of 1 x m.

Superscript T: vector/matrix transpose.

The weight vector ***w*** and bias vector ***b*** are estimated from the training data *D* by using Gradient Descent method^40^ to minimize the overall cost of prediction, or in other words, maximize the probability that ***x*** is in the right category i.

Figure 3.1 shows its architecture.

##### (3) Feed-forward Neural Networks (fully-connected)

The model of Feed-forward Neural Networks has a very similar purpose compared with Multinomial Logistic Regression model. But instead of directly connecting the input ***x*** and output ***y*** using a weight vector ***w*** and a bias vector ***b***, it has several hidden layers of neuron between the input layer and the output layer. A neuron is the simplest processing unit in the model and its output is computed from the following formula^41^:

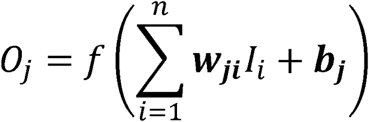

*O*_*j*_: Output of the jth neuron.

*f*: Activation (or transfer) function.

***w***_*ji*_: Synaptic weight corresponding to ith synapse of jth neuron.

***b***_*j*_: Bias of the jth neuron.

*I*_*i*_: ith input signal to the jth neuron.

*n:* Number of input signals to the jth neuron.

An activate function would give more power to the FNNs model compare with the MLM mentioned above. There are many types of activate functions for non-linearizing the models^42^: Conventional sigmoid, such as the Hyperbolic Tangent (f(z) = (exp(z) - exp(-z))/(exp(z) + exp(-z))) and Logistic Function (f(z) = 1/(1 + exp(-z))); Rectified Linear Unit (ReLU) (f(z) = max(0,z)), which has been widely used in recent years. Here we chose the ReLU as the activate function, because it does not require a normalized data input to avoid vanishing gradient problems, so that the model training process would be much more efficient^43^.

Figure 3.2 shows its architecture.

##### (4) Convolutional Neural Networks (locality-sensitive, 2D or 3D)

###### 1. Description of the data manipulation (4×4×4 3D matrix)

In order to use the Convolutional Neural Networks, first we have to convert the shape of input data, from the shape of 1-D vectors (1×64) to the shape of 3-D vectors (4×4×4). Briefly, all the neighboring codons have a 1-bp difference with each other: codon difference of first position was in the depth direction, codon difference of the second position was in the height direction, and the codon difference of the third position was in the width direction (See Figure S3 for details).

###### The classic Convolutional Neural Networks (2D CNNs)

The model of Convolutional Neural Networks (CNNs), which was inspired by visual cortex of the brain, was designed to process multi-dimensional data such as images. Local connectivity, invariance to location and invariance to local transition are three of the advantages it is supposed to have^44^, and the first of which has been appreciated in this study (for detecting the 1-bp change). Typically, there are three different types of layers in the basic structure: convolution layers, pooling layers and fully-connected layers.

For convolution layers, sub-region correlations regardless of their original position in data would be detected. The input (2-D to 3-D vectors or previous layers of feature maps) would be converted to groups of local weighted sums called feature maps, by calculating the convolutions between its local patches and weight vectors called filters^45^. The following formula demonstrates its mathematical process (assuming stride one and one layer zero padding around, modified from the original formula)^46,47^:

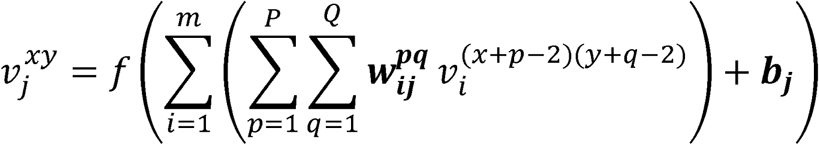

v_*j*_^*xy*^: The output value of an unit at position (x,y) in the jth feature map.

*f*: Activation (or transfer) function.

*m:* Number of layers for the current feature map.

*P:* The height of the kernel.

*Q:* The width of the kernel.

***w*_*ij*_**^***pq***^: The weight value at the position (p,q) at the ith layer of the jth kernel.

***b*_*j*_**. The bias value of the jth kernel.

Using the kernal size of 3×3 and adding one layer of zero paddings around the input data, we can get a DxDxN feature map from the DxDxM input (D denotes the length and width of the input and output feature maps, which were exactly the same in this case, and M, N denotes the depth of the input and output, accordingly). As before, ReLU was also used as the activation function here.

For pooling layers, non-overlapping regions in feature maps had been down-sampled by taking the maximum value of the region (here 2×2 were used), so that local features would be aggregated so as to make up bigger and more complex features.

For fully connected layers, the 2D feature maps were finally converted into a 1D feature vector and then was fed forward into a certain number of categories for classification^19^.

Figure 3.3 shows its architecture.

###### The 3D Convolutional Neural Networks

Since a typical 2D Convolutional Neural Networks (2D CNNs) might lost the information in the direction orthogonal to the 2D plane, we built 3D Convolutional Neural Networks (3D CNNs) despite the cost in computational power and system memories^28^.

For convolution layers, the following formula demonstrates its difference compared with the 2D CNNs (assuming stride one and one layer zero padding around, modified from the original formula)^47,48^:

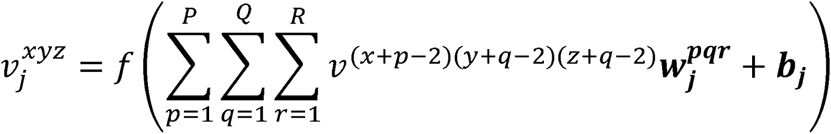

v_*j*_^*xyz*^: The output value of an unit at position (x,y,z) in the jth feature map.

*f*: Activation (or transfer) function.

*P:* The height of the kernel.

*Q:* The width of the kernel.

*R:* The depth of the kernel.

***w*_*j*_**^***pqr***^: The weight value at the position (p,q,r) of the jth kernel.

***b*_*j*_** The bias value of the jth kernel.

Using the kernal size of 3×3×3 and adding one layer of zero paddings around the input data, we can get a DxDxDxN feature map from the DxDxDxM input (D denotes the length, width and depth of the input and output feature maps, which were exactly the same in this case, and M, N denotes the channels of the input and output, accordingly). ReLU was also chosen for activation. For pooling layers, non-overlapping regions now went to 3D (here 2×2×2 were used).

For fully connected layers, similar to 2D CNNs, the 3D feature maps were finally converted into a 1D feature vector and then was fed forward into a certain number of categories for classification.

Figure 3.4 shows its architecture.

**Figure 5.1.**
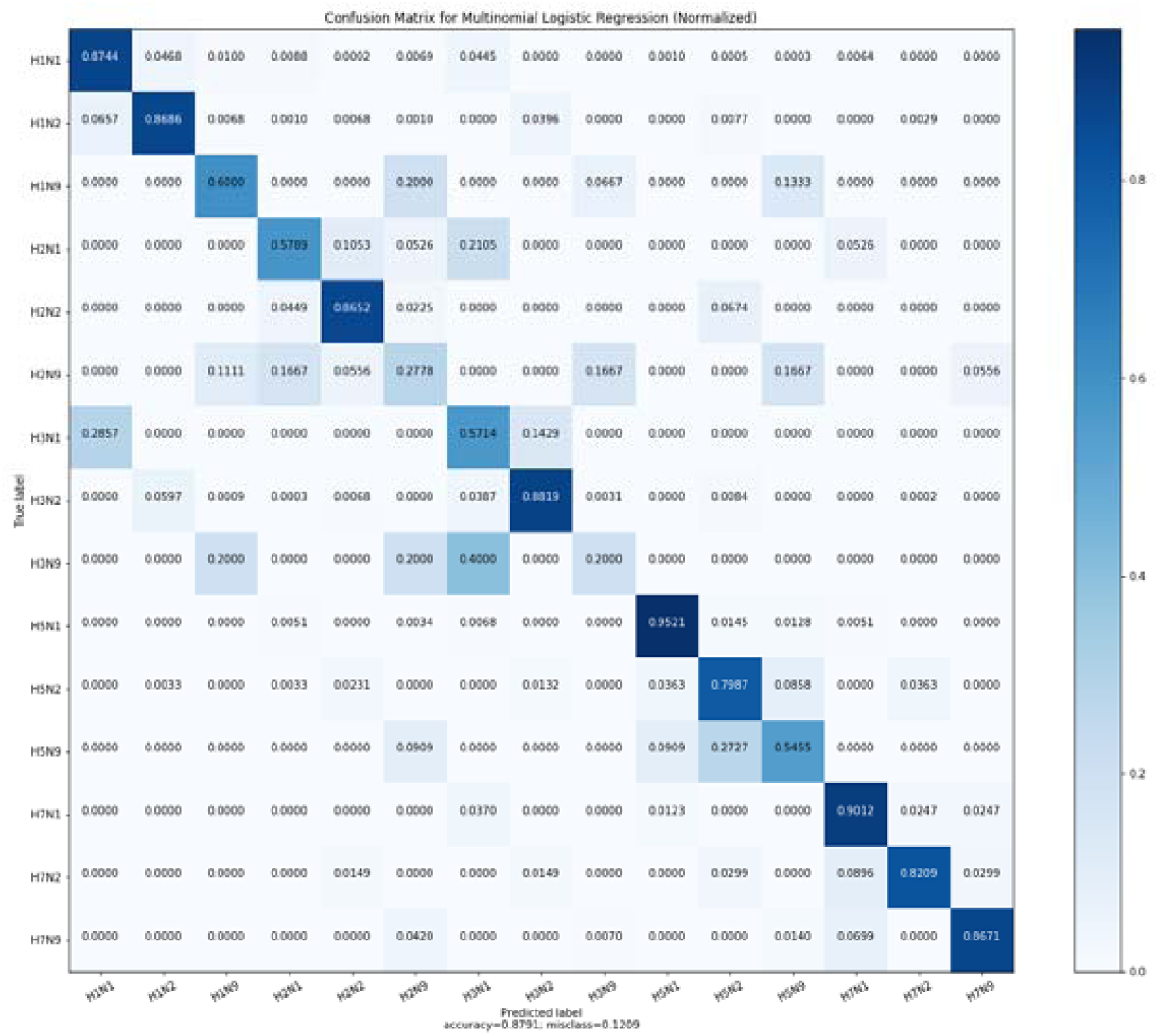
The confusion matrix of Multinomial Logistic Regression model. Normalization according to true label, so the summation of each line is 1.

**Figure 5.2.**
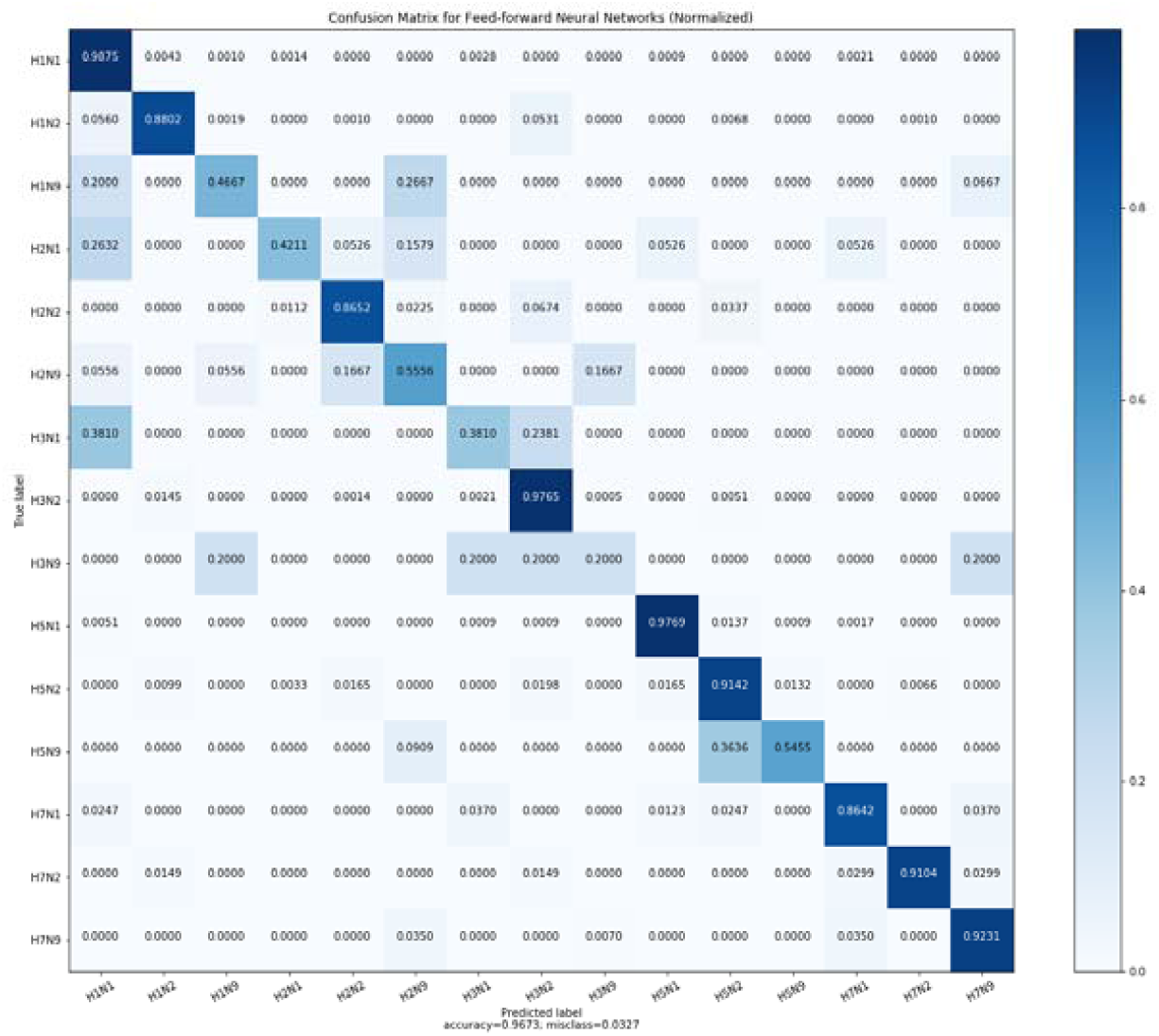
The confusion matrix of Feed-forward Neural Networks model. Normalization according to true label, so the summation of each line is 1.

**Figure 5.3.**
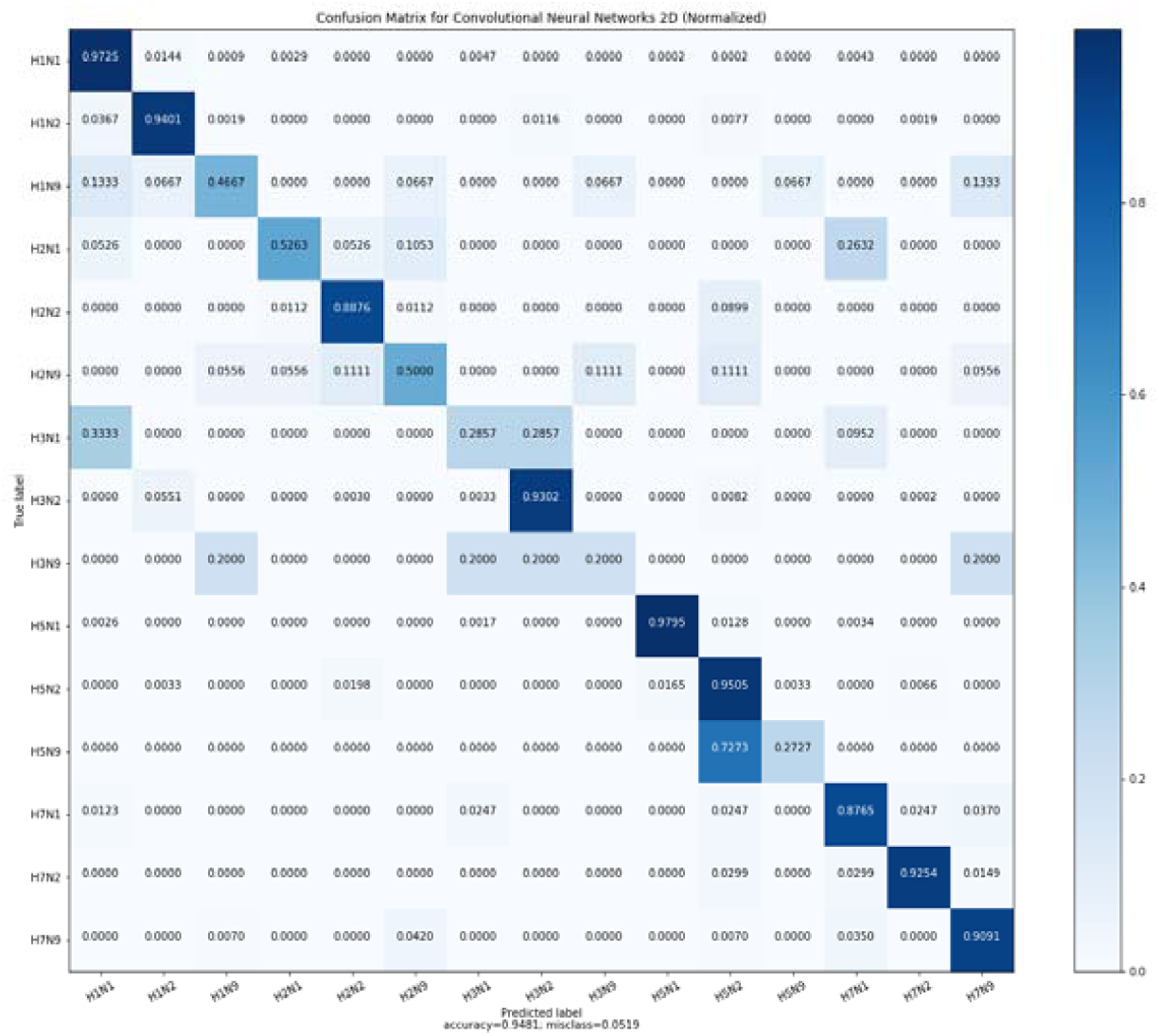
The confusion matrix of 2D Convolutional Neural Networks model. Normalization according to true label, so the summation of each line is 1.

**Figure 5.4.**
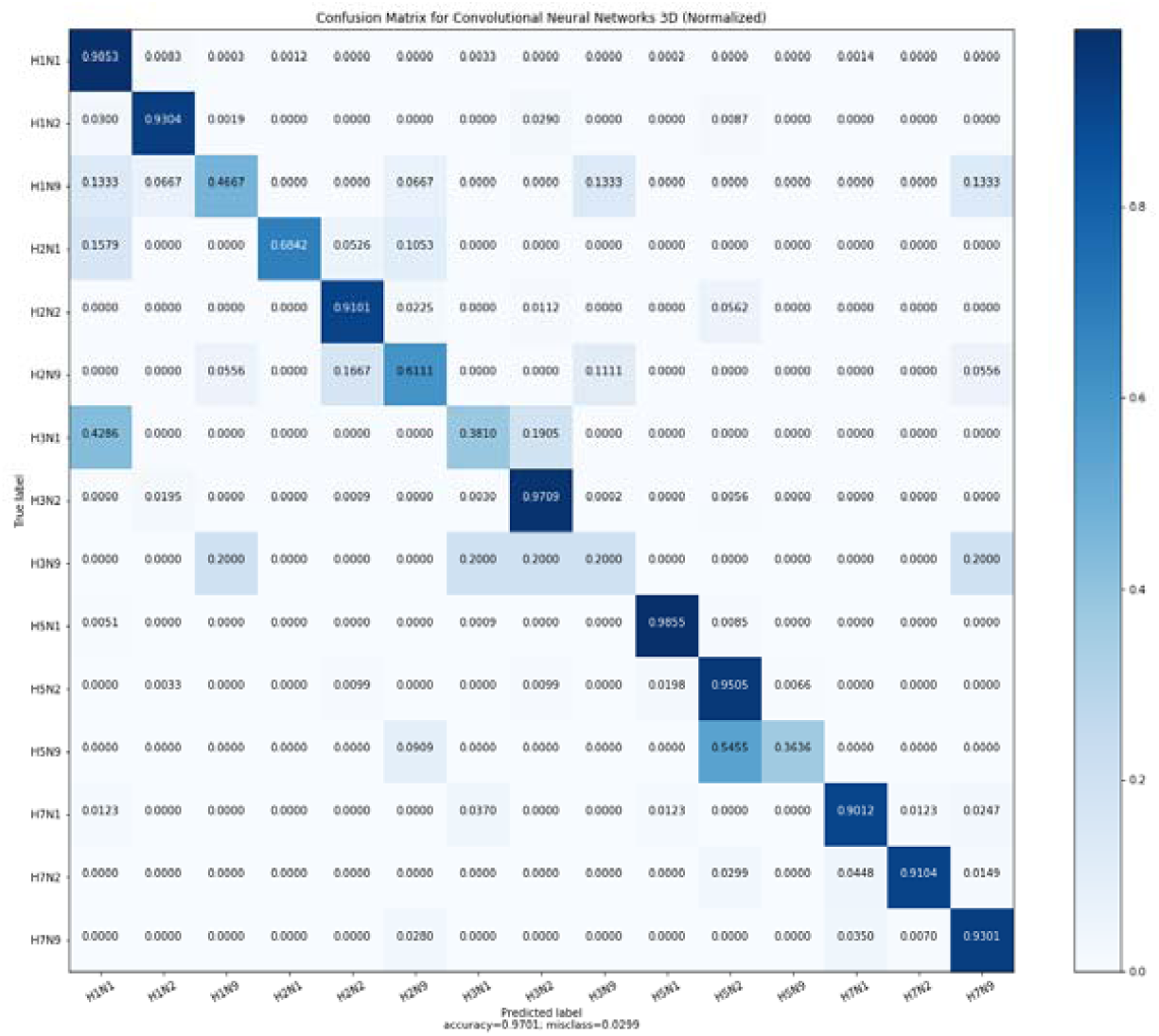
The confusion matrix of 3D Convolutional Neural Networks model. Normalization according to true label, so the summation of each line is 1.

##### (5) Model evaluation

To evaluate the performance of a model, a two-dimensional Confusion Matrix could be calculated^49^. The following parameters could be derived from the matrix^50^:

Recall: TP/(TP + FN).

Precision: TP/(TP + FP).

Accuracy: (TP+TN)/(TP+TN+FP + FN).

F1 score: (2 x Recall x Precision)/(Recall + Precision).

Matthew Correlation Coefficient (MCC):

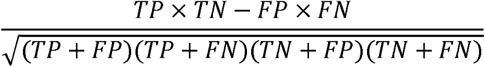

The range is from −1 to +1. A more accurate classifier if the value is towards +1 and the opposite if it is towards-1.

Note: True Positive (TP), False Positive (FP), False Negative (FN), True Negative (TN)

##### (6) Prevent overfitting

Neural networks are so powerful that overfitting could be a very serious problem, thus a very popular method “Dropout” had been applied to prevent this possibility^51^. The keep ratio during each training iteration was 70% in this study.

#### 3) Programming languages and software

##### (1) CodonW software

The computer software CodonW^52^ for calculating the RSCU values mentioned above was downloaded from website (https://sourceforge.net/proiects/codonw/files/codonw/, version 1.4.4).

##### (2) Custom-built scripts and packages

All of the scripts were written in programming language Python (version: 2.7.12). Machine-learning algorithms have been done with the help of the some key packages such as imbalanced-learn (https://pvpi.pvthon.org/pypi/imbalanced-learn. version 0.3.1), scikit-learn (http://scikit-learn.org. version 0.19.1) and TensorFlow (http://www.tensorflow.org. version 1.3.0).

## Supporting information

Benchmarks - continents

Benchmarks - only large samples

Benchmarks

Confusion Matrix

## Additional Figures and Tables

**Figure S1.**
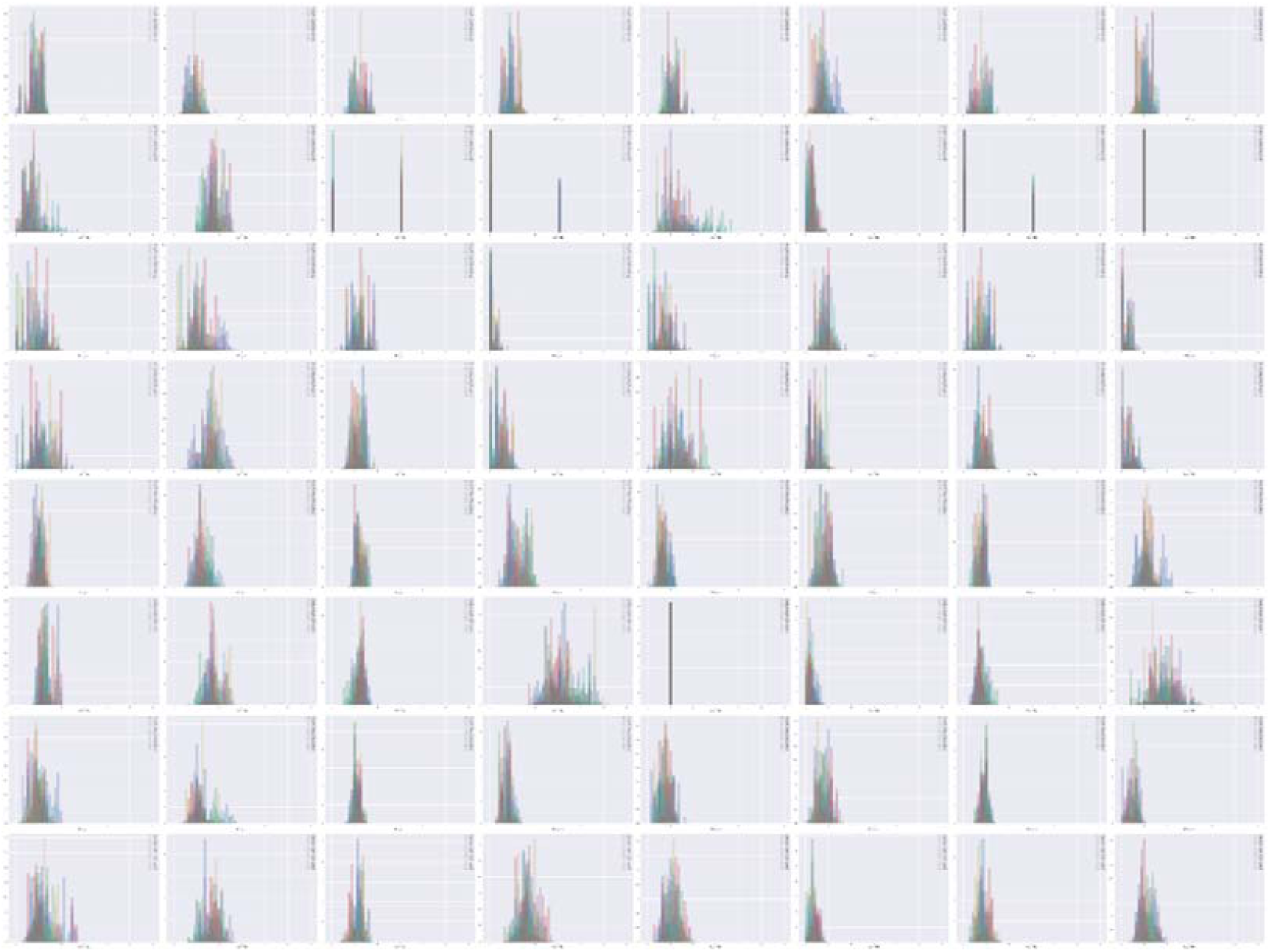
64 RSCUs distributions of different samples. Colors of the bar are used to distinguish the distributions of different serotypes.

**Figure S2.1.**
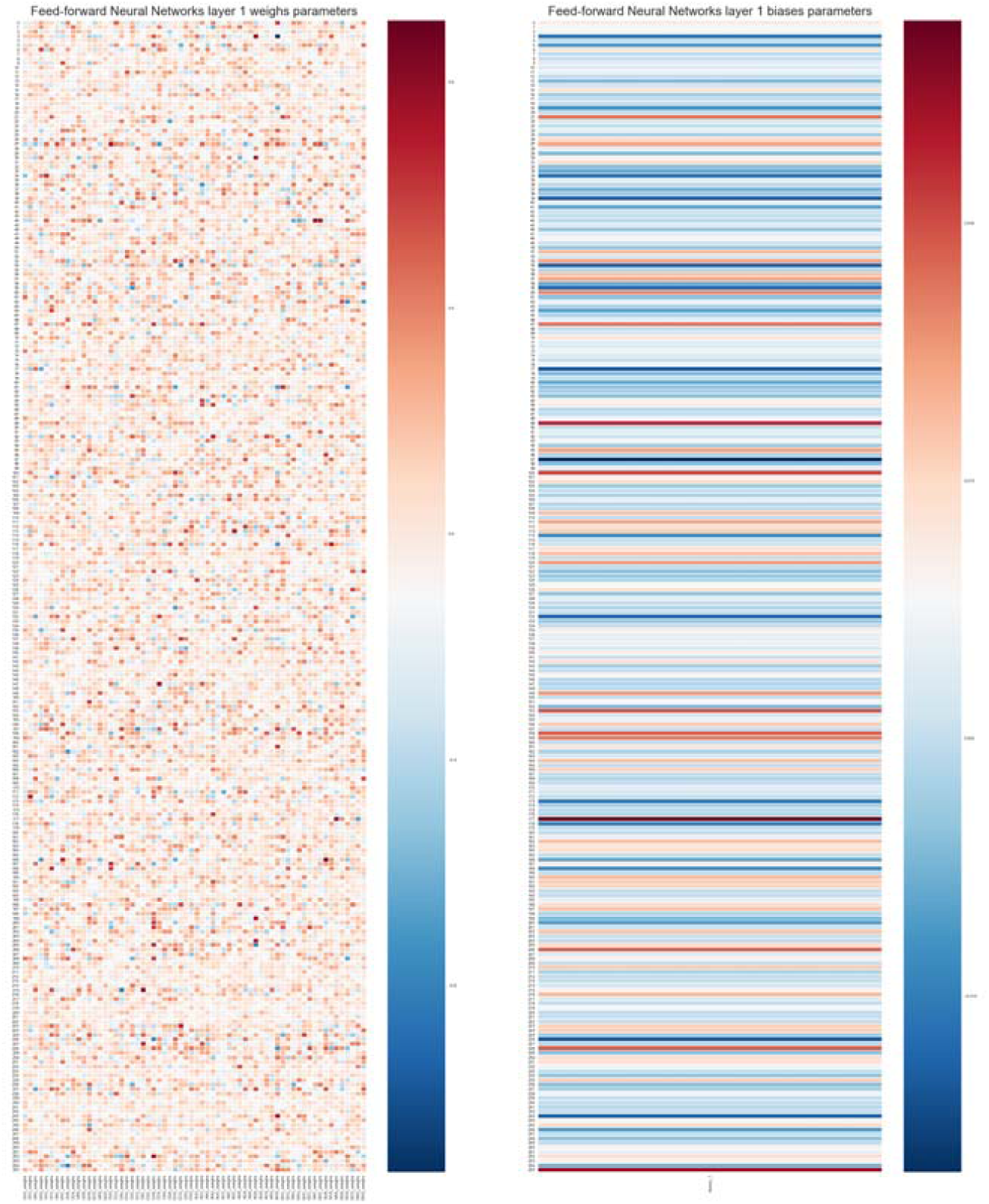
The parameter distribution of Feed-forward Neural Networks model (Layer 1). Left: weighting parameters; Right: bias parameters.

**Figure S2.2.**
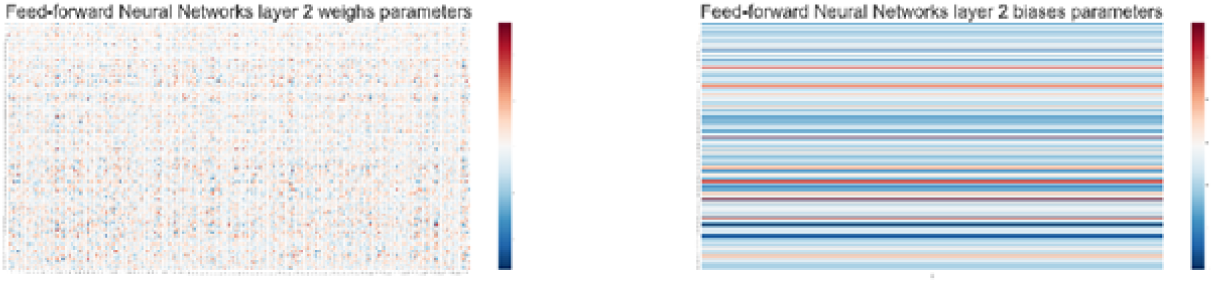
The parameter distribution of Feed-forward Neural Networks model (Layer 2). Left: weighting parameters; Right: bias parameters.

**Figure S2.3.**
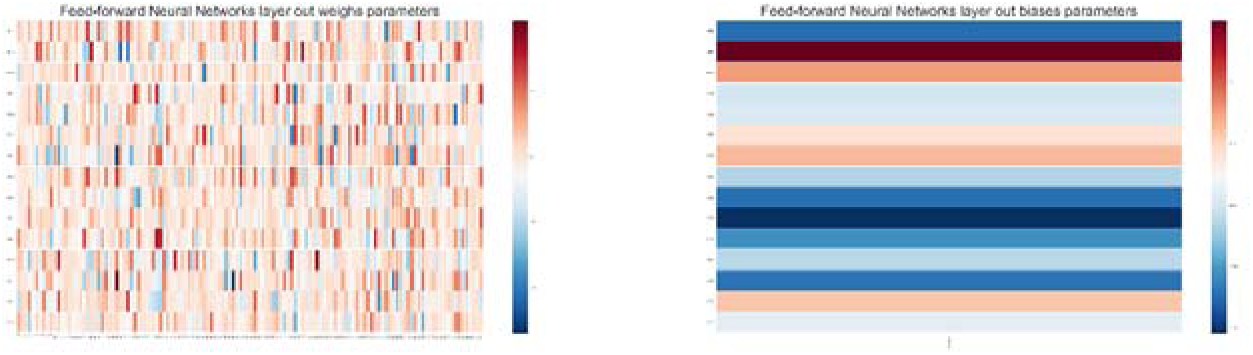
The parameter distribution of Feed-forward Neural Networks model (Layer 3). Left: weighting parameters; Right: bias parameters.

**Figure S3.**
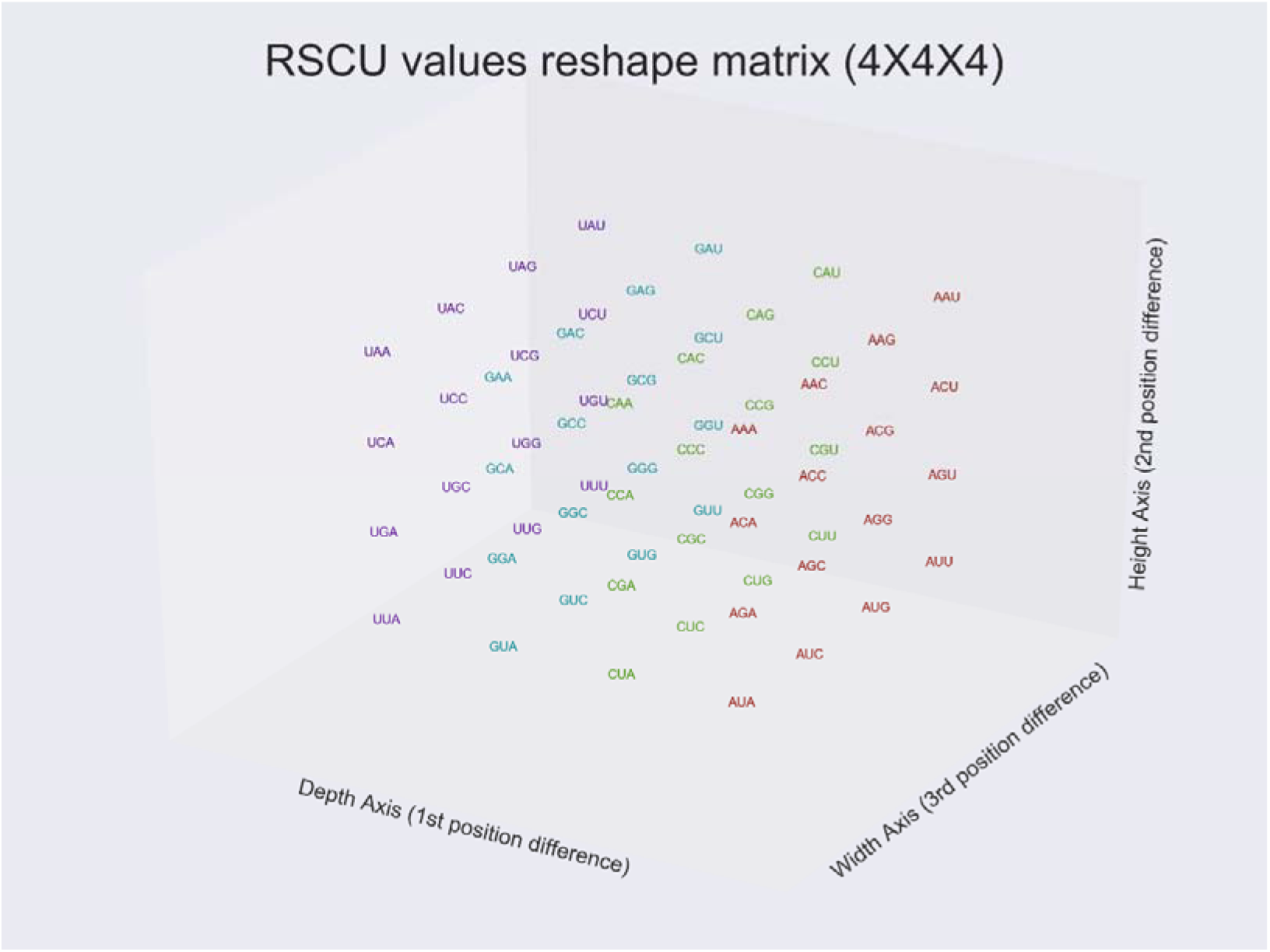
The input 3-D matrix structure after reshaping from the 1-D vector.

**Figure S4.1.**
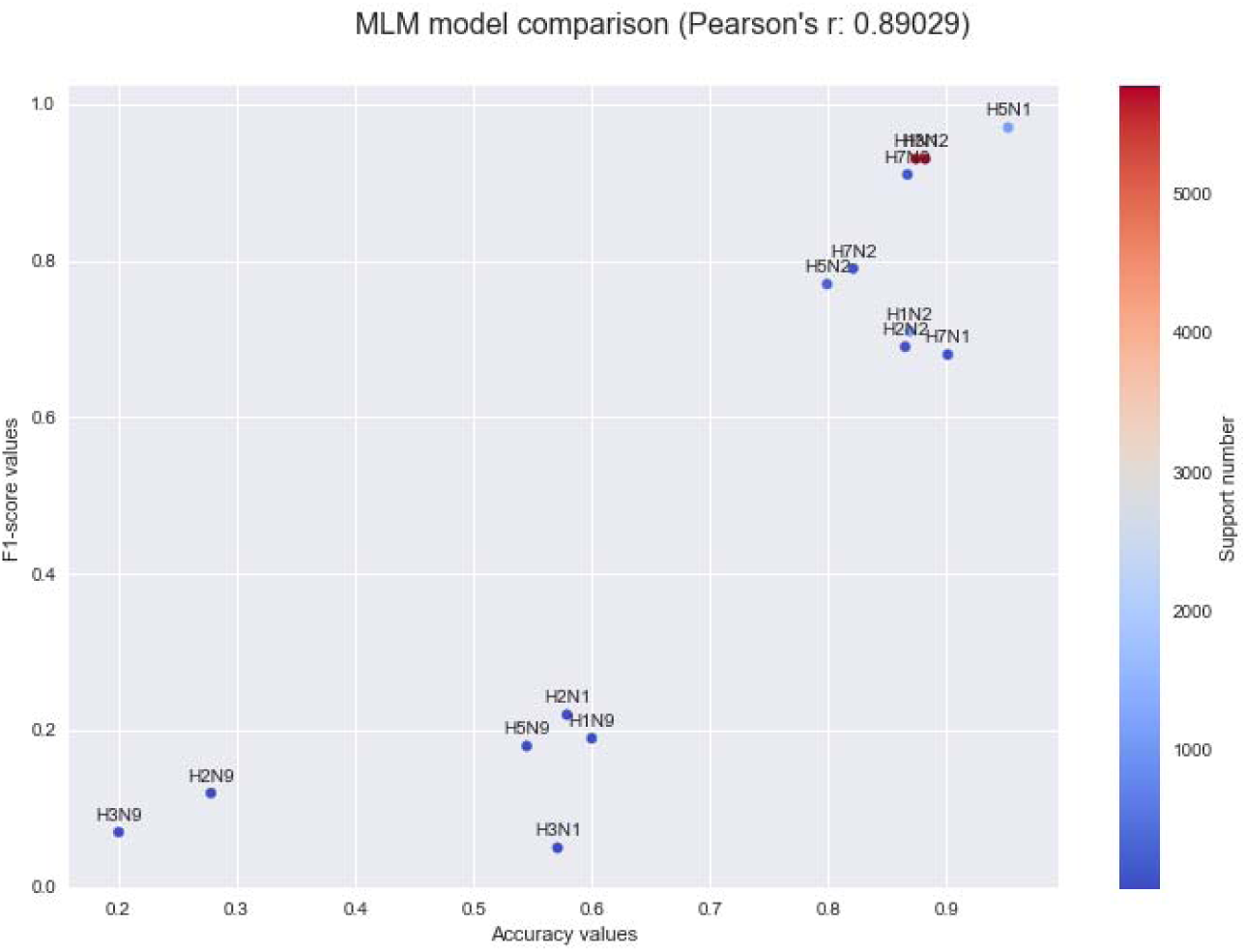
The comparison of Accuracy vs F1 score for the Multinomial Logistic Regression model. Color gradient represents the number of test sample for the serotypes.

**Figure S4.2.**
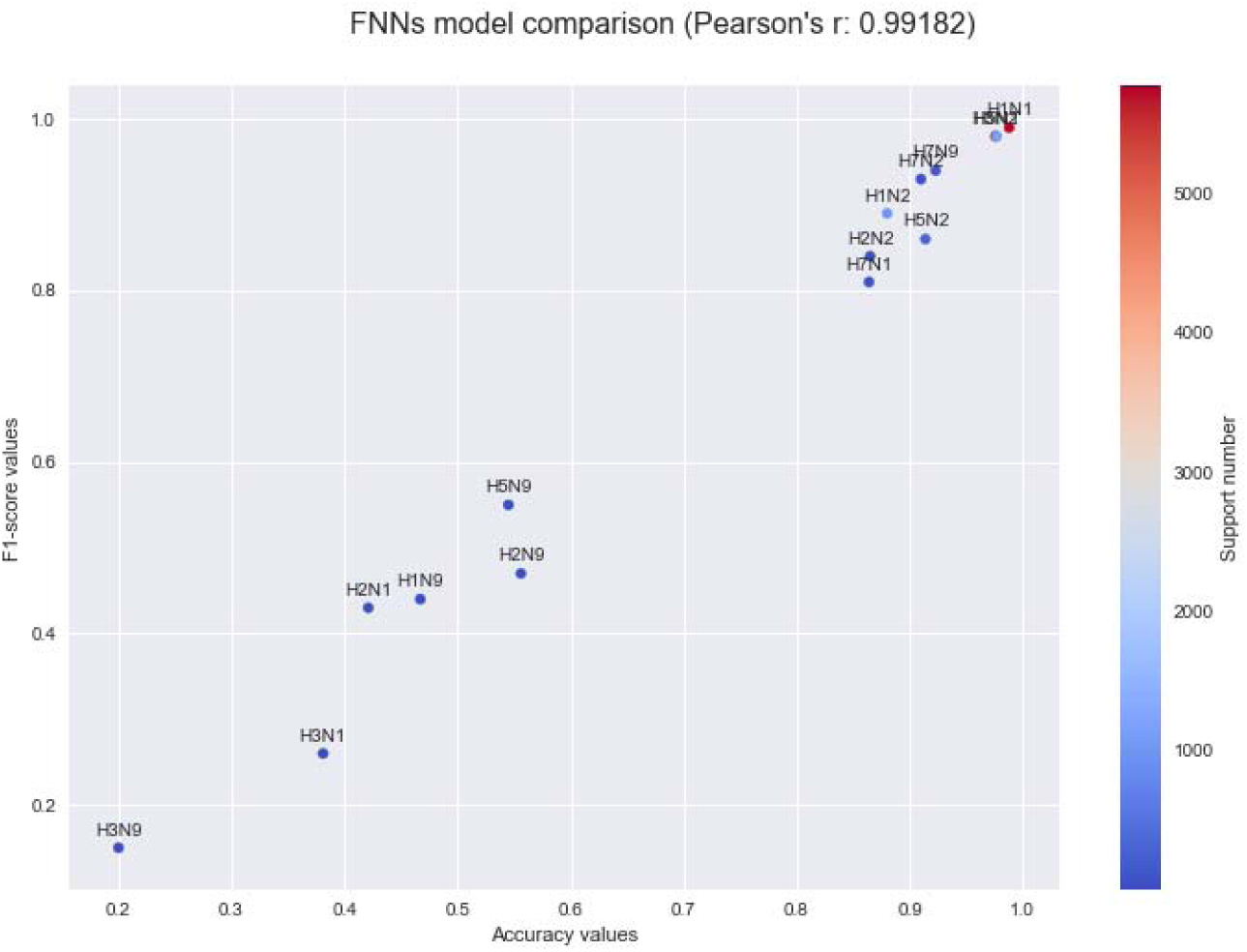
The comparison of Accuracy vs F1 score for the Feed-forward Neural Networks model. Color gradient represents the number of test sample for the serotypes.

**Figure S4.3.**
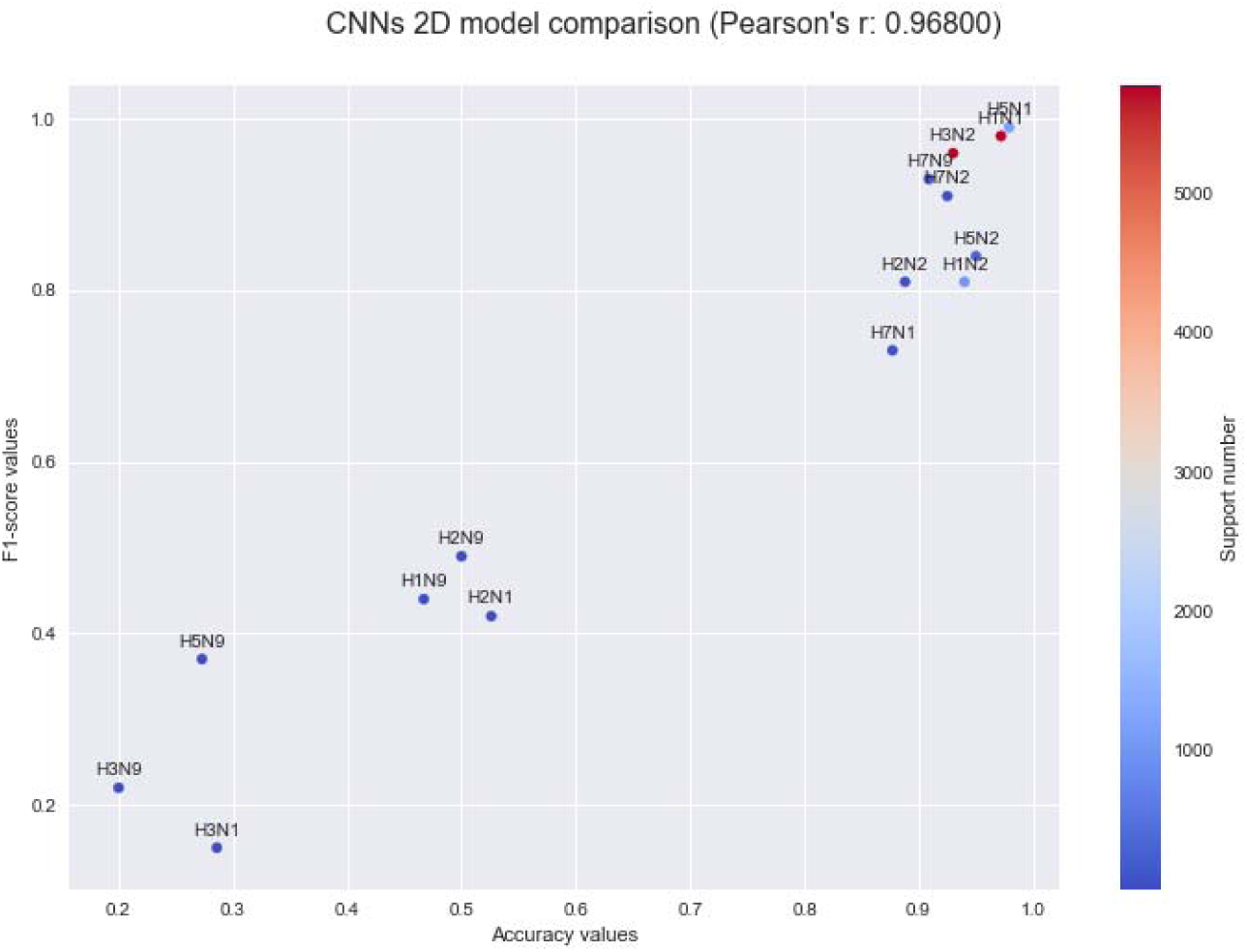
The comparison of Accuracy vs F1 score for the 2D Convolutional Neural Networks model. Color gradient represents the number of test sample for the serotypes.

**Figure S4.4.**
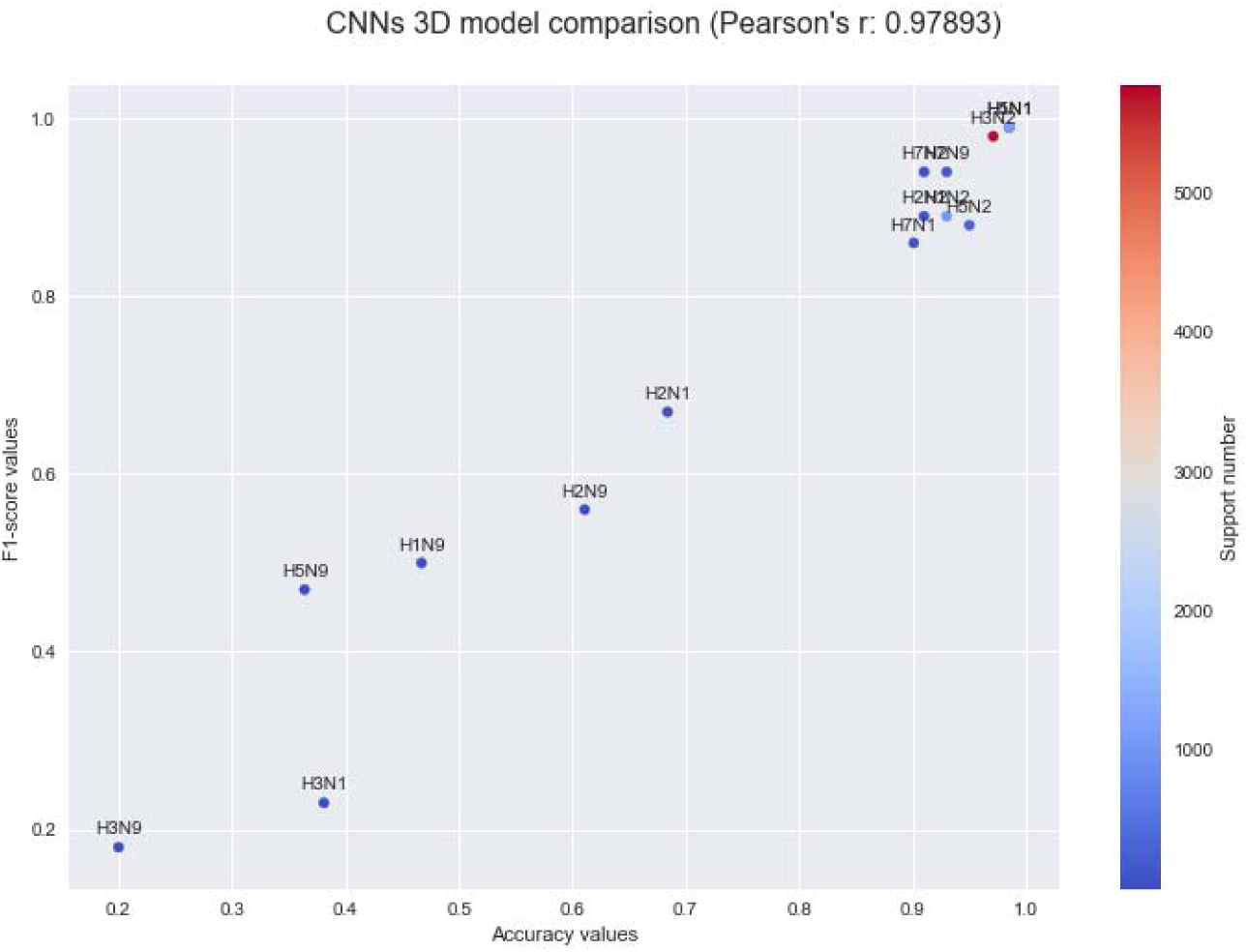
The comparison of Accuracy vs F1 score for the 3D Convolutional Neural Networks model. Color gradient represents the number of test sample for the serotypes.

**Figure S5.1.**
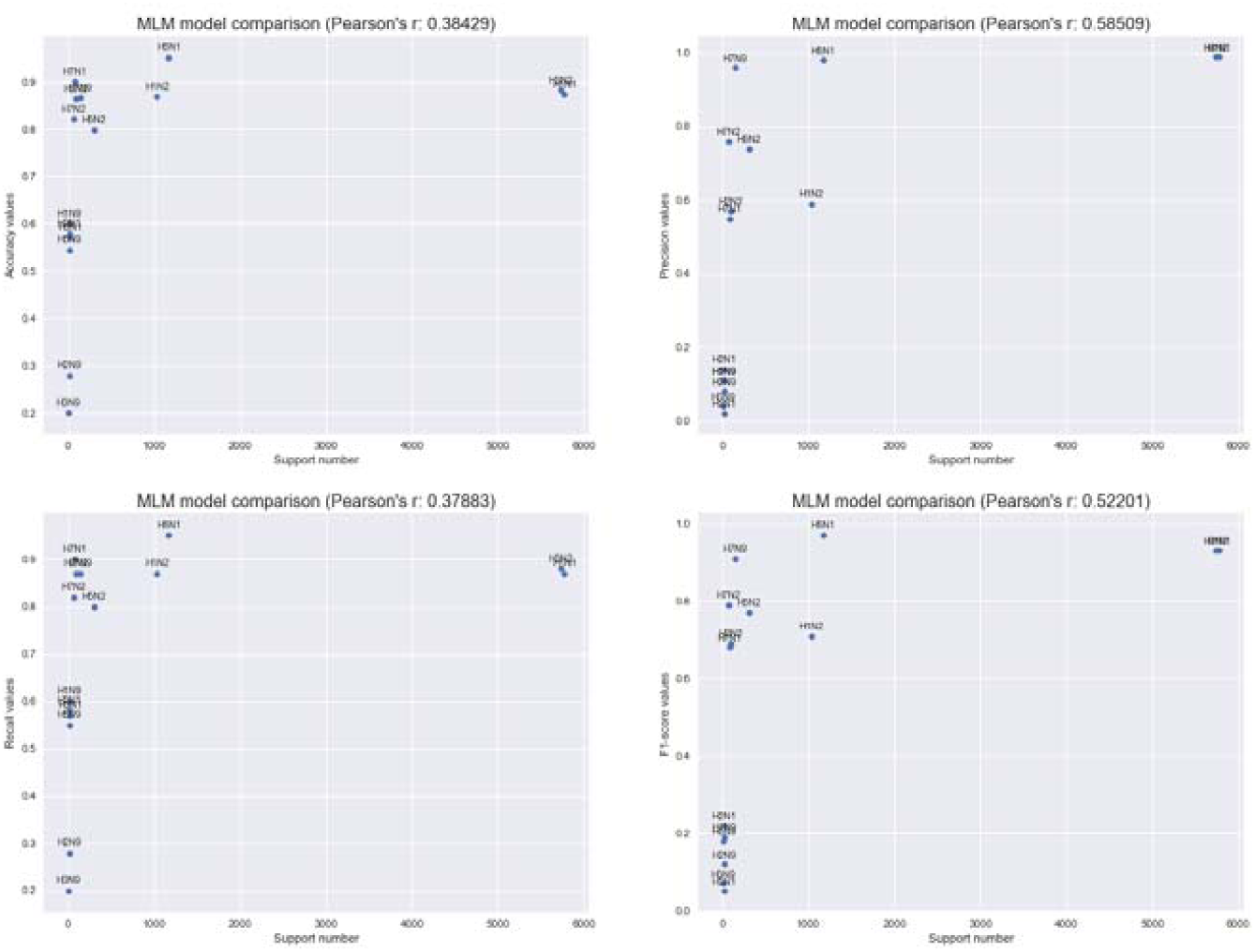
The relationship between test sample support number of each serotype and their evaluation criteria for the Multinomial Logistic Regression model.

**Figure S5.2.**
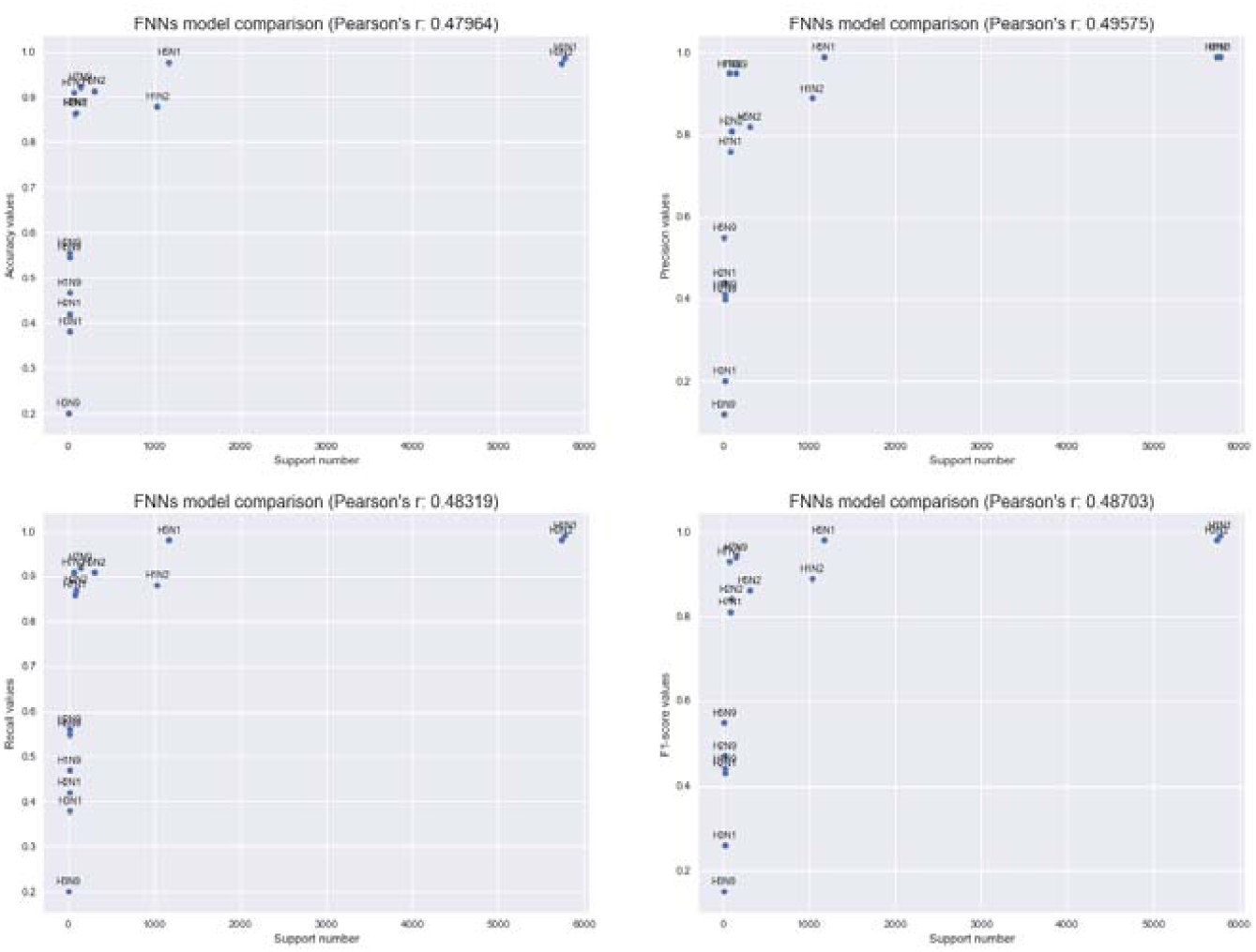
The relationship between test sample support number of each serotype and their evaluation criteria for the Feed-forward Neural Networks model.

**Figure S5.3.**
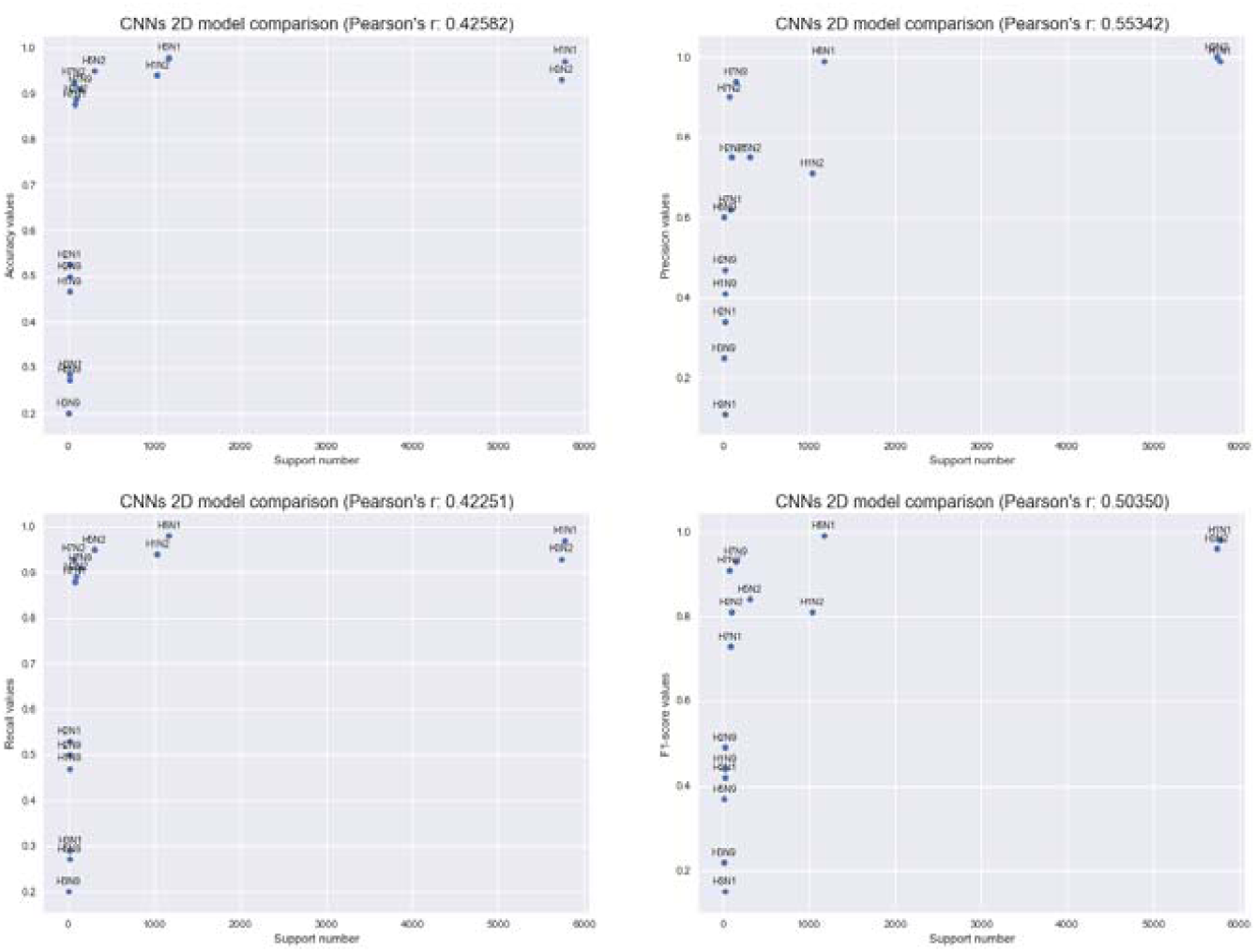
The relationship between test sample support number of each serotype and their evaluation criteria for the 2D Convolutional Neural Networks model.

**Figure S5.4.**
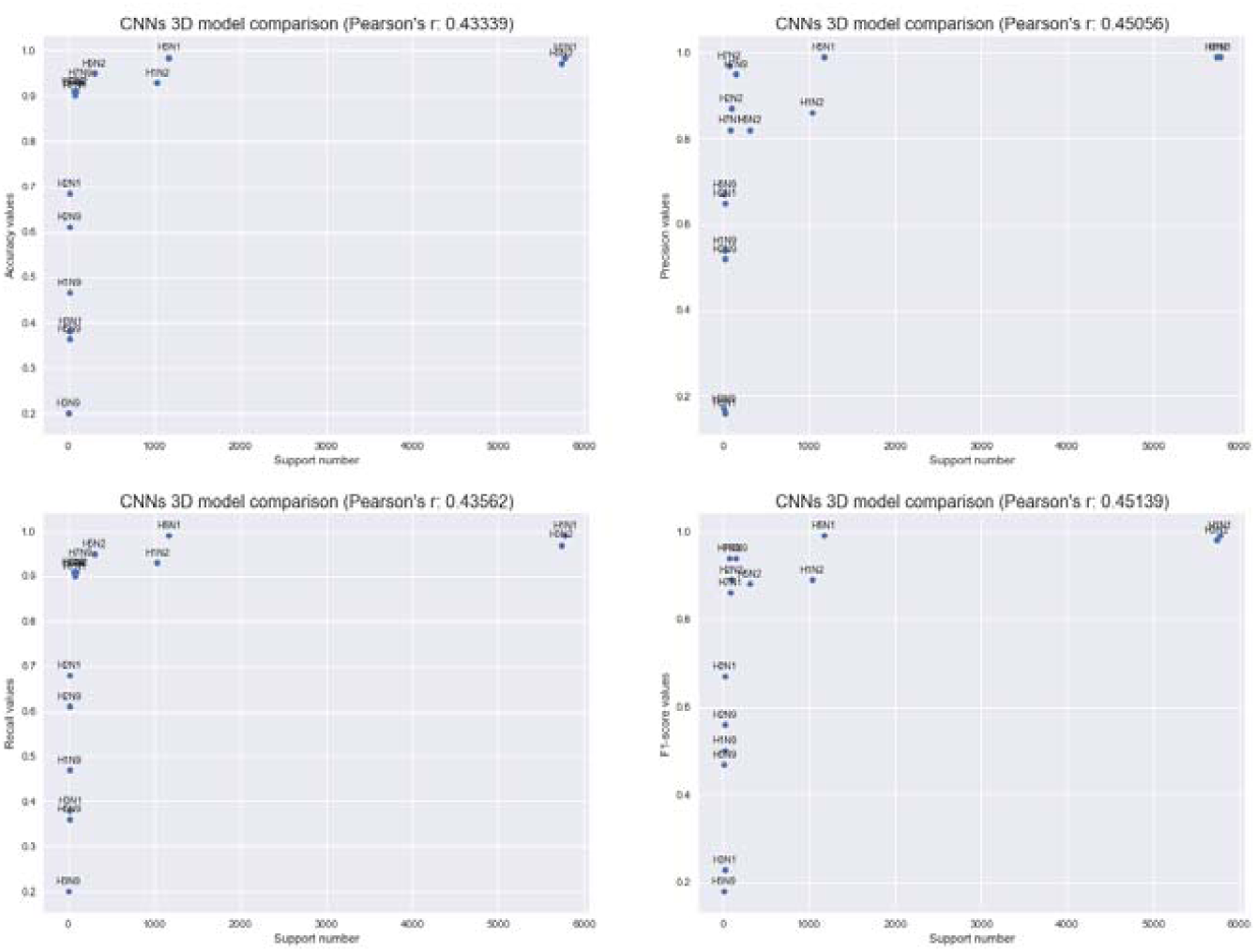
The relationship between test sample support number of each serotype and their evaluation criteria for the 3D Convolutional Neural Networks model.

